# A High-Resolution *N*-Glycoproteome Atlas Reveals Tissue-Specific Glycan Remodeling but Non-random Structural Microheterogeneities

**DOI:** 10.1101/2025.06.24.661310

**Authors:** Yongqi Wu, Yang Muyao, Xu Yongchao, Jia Li, Li Jun, Wang Xiaohan, Zhao Jiayu, Cai Yinli, Zhang Yiwen, Sun Shisheng

## Abstract

The mouse is a widely used model organism in biomedical research, yet a comprehensive understanding of its tissue-specific glycoproteome has been limited due to the structural complexity and microheterogeneity of glycans. Here, we present the most extensive high-resolution *N*-glycoproteomic atlas across 24 mouse tissues, which comprises of 3,045 *N*-glycans with distinct structures attached at 8,681 glycosites on 74,277 glycopeptides and 5,026 glycoproteins. Overall glycan structural patterns show enormous tissue-specific diversities, acting as superior molecular signatures of tissue identity and system origins. Notably, even commonly expressed glycoproteins undergo tissue-dependent glycan remodeling, suggesting that glycosylation may fine-tune protein functions to meet specialized biological demands. These patterns are further shaped by subcellular localization, which constrains glycan variabilities across compartments. Co-occurrence network analyses also expose substructural biases and non-random microheterogeneities among glycans attached at the same glycosites. The dataset serves as a valuable database resource for advancing the structural and functional understanding of glycoproteins.

## Introduction

The mouse is an indispensable animal model in biomedical research^1^. Decades of efforts have mapped the molecular landscapes of mice, including genome^2^, transcriptome^3^, proteome^4^, and phosphoproteome^5^, deepening our understanding of gene regulation and signaling dynamics across tissues. Glycans, another important class of macromolecules, have also shown tissue-specific patterns through previous glycomics studies^6, 7^, which provided an additional dimension to decode the biochemical underpinnings of tissue functions and specialization. However, glycomic studies lack contextual information on protein carriers and specific glycosites, limiting their functional interpretability. Although existing glycoproteomic investigations have also been performed on several individual organs or biofluids, such as liver^8^, brain^9, 10^, and serum^11^, the whole landscape of site-specific glycosylation at the proteome-wide across various tissues remains underexplored.

The microheterogeneity and non-linear structural complexity of *N*-glycans pose considerable analytical challenges. Fortunately, numerous computational tools have been developed for intact glycopeptide analysis over the past decade, including Byonic^12^, GPQuest^13^, pGlyco3^14^, MSFragger-Glyco^15^, O-pair Search^16^, Glyco-Decipher^17^, and GlycanFinder^18^, which are capable of revealing the microheterogeneities of site-specific glycosylation^19^. Nevertheless, most of these tools only identify the predefined glycan compositions, limiting their ability to distinguish functional glycan structures and resolve isomeric glycoforms^12, 13, 20^. Furthermore, the majority of analytical software suffer from the lack of well-established glycan databases for most species as they adopt a glycan database-dependent strategy. To address these limitations, we previously developed a modular strategy that determines detailed structural features of *N*-glycans at specific glycosites by leveraging characteristic B- and Y-ions in tandem mass spectra (MS/MS) acquired with low HCD collision energy^21^. The StrucGP software, based on this strategy, enables the high-throughput de novo structural interpretation of site-specific *N*-glycans^21^, achieving high-resolution glycoproteomic analyses across diverse biomedical samples.

In this study, we generated the first structural and site-specific *N*-glycoproteome atlas across 24 mouse tissues using the StrucGP software. This comprehensive high-resolution dataset provides an unprecedented insight into the tissue-specific landscape of protein *N*-glycosylation, revealing how distinct glycan structures and site-specific glycosylation contribute to tissue identity and specialization. We identified molecular signatures defined by unique glycopeptides, which robustly distinguish tissue types and offer a new dimension for understanding cell and organ differentiation beyond gene and protein expressions. Importantly, we demonstrate that even commonly expressed proteins across various tissues undergo tissue-dependent *N*-glycan remodeling, suggesting that glycosylation fine-tunes protein function in response to specific physiological contexts. Compartment-specific glycosylation patterns further highlight the alignment between glycan features and biosynthetic pathways or functional demands. Through glycan co-occurrence network analyses, we also reveal highly organized, non-random patterns of glycan usage that illuminate the organizational logic of glycoproteome diversity. Collectively, this study not only expands the scope of glycoproteomics but also establishes a foundational resource for functional and mechanistic investigations of glycosylation in tissue biology and mammalian physiology.

## Results

### 1. A High-resolution *N*-Glycoproteome Atlas Across 24 Mouse Tissues

In this study, we created a comprehensive atlas of the mouse *N*-glycoproteome across 24 distinct tissues, encompassing 16 major organs, 7 brain regions, and the serum, collectively representing all eight major anatomical systems. Seven brain regions included the medulla, cerebellum, white matter, hypothalamus, cortex, hippocampus, and olfactory bulb (**Figure 1A**). For each tissue, equal amounts of proteins were pooled from five male and five female C57BL/6N mice (8-9 weeks) to generate 24 tissue-specific samples. Glycopeptides were enriched using hydrophilic interaction chromatography (HILIC) followed by mixed anion-exchange columns (MAX)^22^ and subjected to at least triplicate LC-MS/MS analyses per sample. For 22 tissues (excluding gallbladder and ovary due to limited sample size), the glycopeptides enriched by HILIC were further separated into six fractions and analyzed by at least duplicate LC-MS/MS runs per fraction. These resulted in a total of 456 raw data (**Figure 1B**). During mass spectrometry analyses, each glycopeptide peak was first fragmented by a relatively high HCD energy (HCD=33%) followed by a relatively low HCD energy (HCD = 20%) for peptide sequence and glycan structure analyses, respectively^21^.

**Figure 1.**
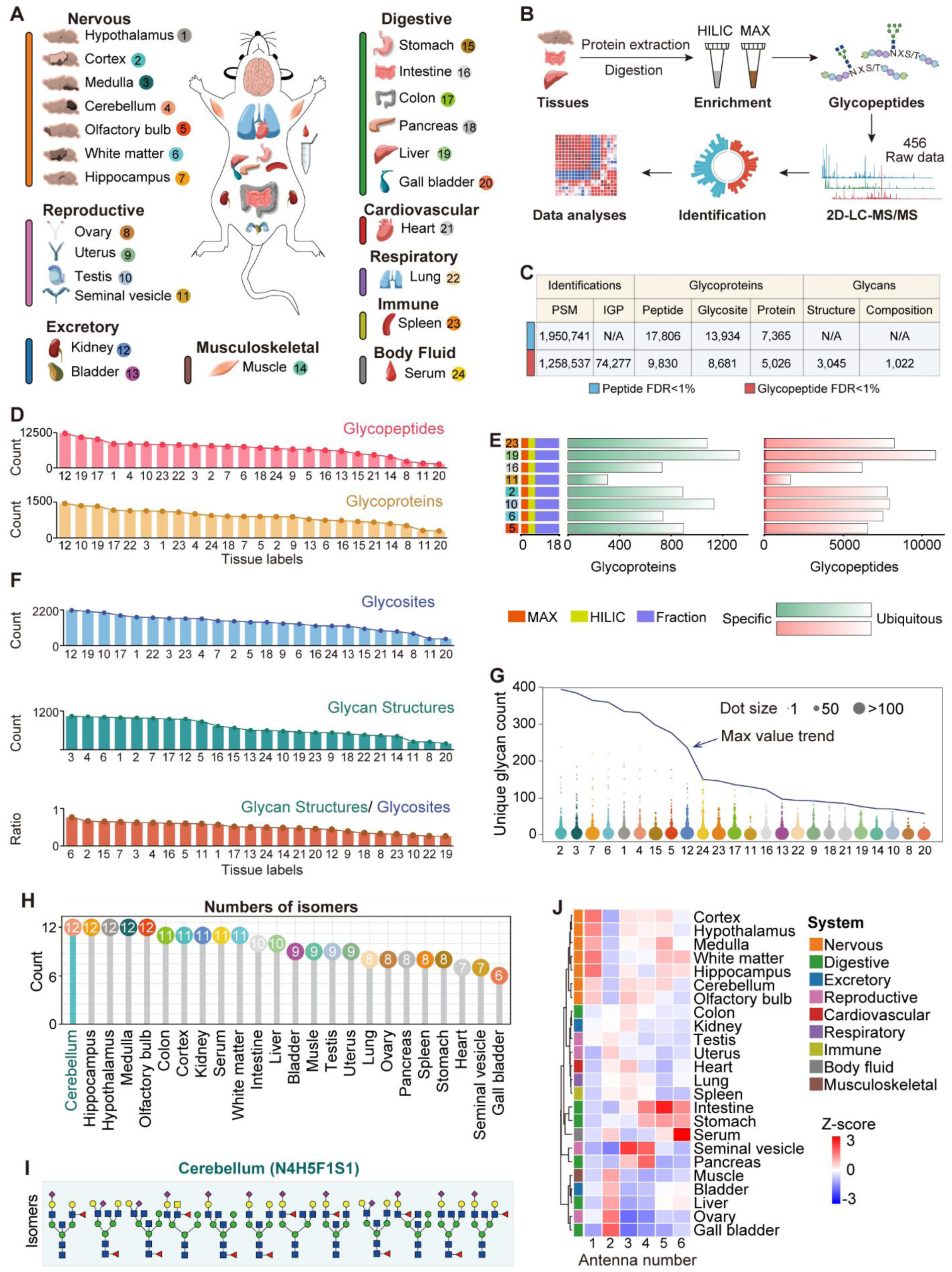
Comprehensive mapping of high-resolution *N*-glycoproteome across 24 mouse tissues. (A) Schematic of the study design, depicting structural and site-specific glycoproteome across 24 tissues (covering 8 major systems) from C57BL/6N mice (5 males and 5 females, 8–9 weeks). Each tissue is labeled by a number, and each organ system is represented by a unique color code. (B) Flowchart overview of generating mouse multi-tissues *N-*glycoproteome. The whole process includes tissue collection, protein extraction and digestion, glycopeptide enrichment, mass spectrometry analyses, database search for glycopeptide identification, and data analyses. (C) Intact *N*-glycopeptides identified based on FDR <1% at the glycosite-containing peptide level (upper) or at both glycosite-containing peptide and glycan levels (lower) (see Table S1). PSMs: peptide-spectrum matches; IGPs: intact glycopeptides. (D) Total number of intact glycopeptides (upper) and glycoproteins (lower) identified from each tissue. (E) Comparison of glycoprotein and glycopeptide counts identified from eight tissues with equal LC-MS runs. (F) Identification of glycosites, glycan structures and the ratio of glycans per glycosite across tissues. (G) Number of glycan structures identified at individual glycosites across 24 tissues. The maximum number per site is highlighted by a line overlay. (H) The highest number of glycan structure isomers identified from each tissue. (I) Twelve different glycan structures discriminated from glycan composition of N5H5F1S1 in cerebellum (see also Figure S4A). (J) Distribution of different antenna counts in site-specific glycans identified from 24 mouse tissues (see also Figure S5).

Using StrucGP, we identified 17,806 glycosite-containing peptides from 1,950,741 oxonium ion containing spectra, representing 13,934 *N*-glycosites on 7,365 *N*-glycoproteins from 24 mouse tissues within 1% false discovery rate (FDR) at the peptide level (regardless of their attached glycans) (**Figure 1C**). After further filtering by FDR<1% at the glycan level, a total of 74,277 unique intact *N*-glycopeptides (IGPs) were identified from 1,258,537 high-quality MS/MS spectra, comprising 3,045 *N*-glycans with distinct structural features and 8,681 *N*-glycosites from 5,026 *N*-glycoproteins (**Table S1**; **Figure 1C**), constituting one of the most extensive tissue-specific glycoproteomic resources to date. Notably, this dataset encompassed 2,582 of the 4,065 *N*-glycoproteins reviewed in UniProtKB (63.5%) and delineated 4,778 previously unannotated glycoproteins, including 2,897 novel ones at the glycan level (**Figure S1A**). It also covered 47.2% of known *N*-glycans (706 compositions) in GlyConnect^23^ and identified 689 novel glycan compositions (**Table S2**; **Figure S1B**), significantly expanding the mouse glycome and glycoproteome. It should be noticed that these novel compositions only represent new combinations of known core and branch structures. Further comparisons with glycan databases supported by software tools such as Glyco-Decipher, Byonic, and pGlyco3 revealed 207 glycan compositions exclusively identified in our results (**Table S2**; **Figure S1C)**. Motif analysis confirmed a higher prevalence of the N-X-T motif over N-X-S as reported previously^24, 25, 26^ (**Figure S1D**). Across the 24 mouse tissues, glycoprotein and glycopeptide identifications varied substantially across tissues. The gallbladder yielded 1,304 unique *N*-linked IGPs from 292 glycoproteins, while 12,208 IGPs form 1,441 glycoproteins derived from the kidney. This disparity is likely arising from the absence of fractionation in the gallbladder samples, highlighting the critical role of sample fractionation in enhancing the depth of analysis for complex tissues (**Figure 1D**). Notably, even with an equal number of LC-MS/MS runs, the number of identified glycoproteins and glycopeptides varied significantly across tissues^27^. As shown in **Figure 1E**, the liver identified the most glycoproteins and glycopeptides, whereas the seminal vesicles identified the fewest, which is consistent with previous protein identifications from multi-tissue proteomics^28^. Besides, replicate analyses of the same fraction revealed high correlation coefficients at both glycan (R=0.59–0.88) and glycosite (R=0.61–0.83) levels (**Figure S2A**), and the inter-tissue variability greatly exceeded technical variation, further confirming the robustness of the dataset (**Figure S2B**).

We further profiled tissue-specific *N*-glycosylation landscapes and uncovered pronounced differences in glycan diversity and complexity across organs. By quantifying the number of identified *N*-glycans per tissue and analyzing glycosite-specific occupancy, we found that the kidney harbored the most glycosites (2,181 glycosites modified with 952 unique glycans) at both peptide and glycopeptide levels (FDR <1%), likely due to more extensive mass spectrometry coverage (**Figures S3A** and **S3B**). In contrast, nervous system tissues (seven brain regions) showed relatively fewer glycosites (1,337-1,755) but featured the greatest structural diversity, with 869-1,042 unique glycans identified across brain regions. These patterns were further illustrated by the glycan-to-glycosite ratios: the kidney showed a relatively low ratio of 0.44, while the seven brain regions ranged from 0.57-0.76; notably, the liver exhibited the lowest ratio at 0.26 (**Figure 1F**). In neural tissues, individual glycosites carried up to 275-393 distinct glycan structures, highlighting exceptional microheterogeneitie compared with other tissues (**Figure 1G**). In addition, the diversity of *N*-glycans in neural tissues was further evidenced by the larger number of structural isomers identified for individual glycan compositions. Isomeric analysis revealed that each glycan compositions could be further expanded into 6-12 isomers across tissue, with the nervous system containing the most (**Figure 1H**). These structural isomers were distinguished by characteristic B and Y ions in low HCD energy of MS/MS, as exemplified by the 12 isomers of the composition N5H5F1S1 (**Figures 1I** and **S4A**), and in many cases also resolved by C18 chromatography separation based on distinct retention time (**Figure S4B**). Despite the remarkable structural diversity, *N*-glycans in the nervous system tended to exhibit relatively low complexity, as the glycans in nervous system was predominantly mono-antennary, consistent with previous finding in neuronal tissues^29^. In contrast, bi- and tri-antennary glycans dominated most tissues, while tri- and tetra-antennary glycans were especially abundant in exocrine organs like the pancreas and seminal vesicles. Notably, highly branched glycans were observed in the stomach, intestine, and serum (**Figures 1J** and **S5**).

## 2. Site-Specific Glycans Show Tissue-Specific Structural Features

To investigate tissue-specific glycan preferences, we first extracted the top ten most abundant *N*-glycans in each tissue based on the numbers of their modified glycosites, excluding commonly high-abundant oligo-mannose glycans. The nervous system and kidney were enriched in core-fucosylated bisecting glycans (N3H3F1). In contrast, although both kidney and seminal vesicle exhibited high level of Lewis^x/a^ and Lewis^y/b^ structures, bisecting glycans were far less abundant in the seminal vesicle. Liver and serum were characterized by complex, sialylated glycans with relatively low levels of core fucosylation. Testis-specific glycans were notably enriched in terminal GlcNAc. In addition, the pancreas, seminal vesicle, and several digestive tissues—such as the intestine, colon, and stomach— shared an enrichment of terminal galactose. Despite this common feature, the most abundant glycans in pancreas were hybrid type, while digestive tissues predominantly carried bisected core structures. The seminal vesicle, by contrast, was distinguished by strong fucosylation on terminal galactosylated glycans, lacking the bisecting core features observed in the digestive tract. (**Figure 2A**).

**Figure 2.**
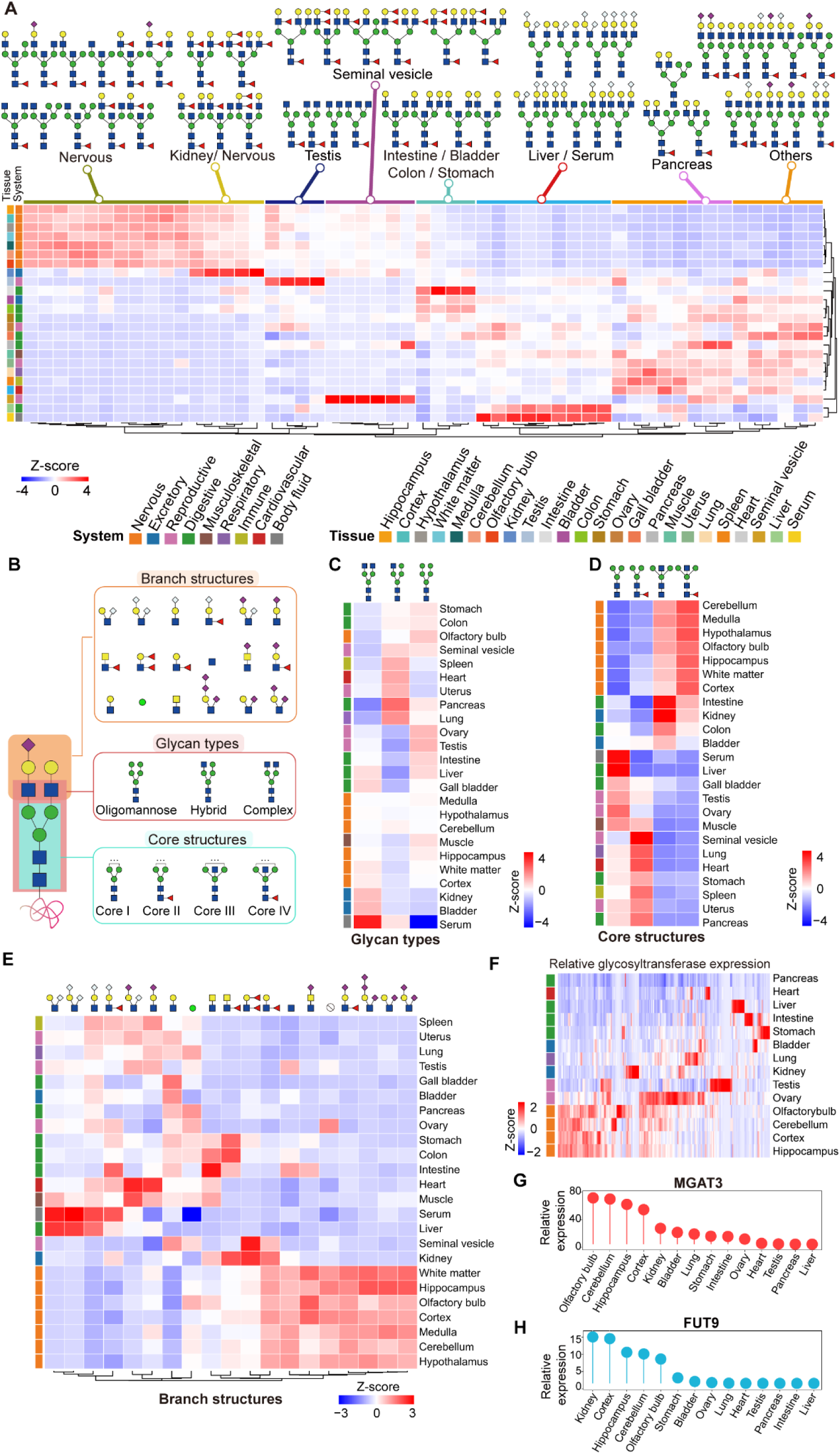
Distinct glycan structural patterns and substructural features across 24 mouse tissues. (A) Heatmap shows top 10 glycan structures identified in each tissue ranked by the numbers of modified glycosites. (B) Classification of 3 glycan types, 4 core structures (Core I–IV), and 17 branch structures across mouse tissues based on structural modules. (C-E) Distribution of different modules of site-specific *N*-glycans identified from 24 mouse tissues, including three glycan types (C), four core structures (D), and 17 branch structures (E). The percentages were calculated based on unique glycopeptides containing each type of glycan structural feature versus total glycopeptides identified from the tissue. (F) Heatmap showing mRNA expression levels of 209 glycosyltransferases across 14 tissues using previously published RNA-seq data (see Table S4). (G and H) Relative expression levels of glycosyltransferases (Mgat3, Fut9) across 14 mouse tissues based on RNA-seq data. Mgat3, Beta-1,4-mannosyl-glycoprotein 4-beta-N-acetylglucosaminyltransferase. Fut9, 4-galactosyl-N-acetylglucosaminide 3-alpha-L-fucosyltransferase 9.

Based on the modular structures of identified *N*-glycans across mouse tissues, three glycan subtypes with four core structures and 17 branch structures were observed (**Table S3**; **Figure 2B**). Among glycan subtypes, serum glycoproteins were predominantly modified with complex glycans (74.9% IGPs), with minimal oligo-mannose glycans (3.1% IGPs). Complex glycans were also attached at more than 50% IGPs in bladder, kidney, gall bladder, liver, white matter, contex, hippocampus, and hypothalamus. Hybrid glycans were highly expressed in the pancreas (30.7%), lung, heart, spleen, and uterus (25.3%); whereas oligo-mannose glycans were primarily detected in the testis (38.1%), ovary, olfactory bulb, intestine, liver (34.2%), etc (**Figure 2C**). Among four core structures, typical core structure (HexNAc_2_Hex_3_, Core I) was the most prevalent, showing notably higher abundance in serum (73% vs. 48.5% across all tissues) and liver (74.3%). The fucosylated core structure (Core II) was enriched in the seminal vesicle (51.9% vs. 33.6% across all tissues), heart (46.1%), and lung (45.2%). The bisected core structure (Core III), besides being enriched in the nervous system (averaged 7.2% in brain regions vs. 4.8% across all tissues), was also prominent in digestive and exocrine organs such as the kidney (12.8%), intestine (12.1%), and colon (8.7%). In contrast, the fucosylated bisecting core (Core IV) was specifically elevated in neural tissues, averaging 29.4% across brain regions compared to 13.1% in all tissues (**Figure 2D**). Interestingly, more pronounced tissue-specific distributions emerged when examining distinct glycan branch structures. The nervous system predominantly featured terminal GlcNAc^30^, unbranched structures (labeled as Ø), Lewis^x/a^, and various sialylated branches (except sialyl-LacNAc) mainly modified by N-acetylneuraminic acid (Neu5Ac) but lack the N-glycosylneuraminic acid (Neu5Gc), in line with previous reports^31^. Conversely, serum and liver glycans displayed a predominance of Neu5Gc-containing branches in the absence of Neu5Ac^32^. Heart and muscle showed a relatively high abundance of Sialyl-LacNAc via Neu5Ac, or with an additionally branched Neu5Gc. Additionally, rare branch structures such as LacdiNAc-containing glycans (excluding sialyl-LacdiNAc which tended to cluster with other sialylated branch structures in neural tissues) were selectively enriched in digestive organs including the stomach, colon, and intestine, while branches with antenna-fucosylation (without sialylation) were highly expressed in kidney, and the Lewis^y/b^ epitope was also highly expressed in the seminal vesicle (**Figure 2E**).

Given that glycosyltransferases (GTs) are key enzymes governing *N*-glycans biosynthesis by catalyzing glycosyl transfer reactions^33^, we further explored their expression patterns. Due to limited GT identification in proteomic data, we analyzed previously reported RNA sequencing data across multiple mouse tissues^4^, successfully matching 10 peripheral tissues and 4 brain regions with our study samples. The comparative analyses identified 209 GTs displaying distinct tissue-specific expression profiles (**Table S4**; **Figure 2F**). Notably, the four brain regions exhibited a unique GT expression pattern, consistent with the specific distribution patterns of *N*-glycans observed in our study. As expected, the elevated levels of Core III and Core IV in the brain regions strongly correlated with high expression of the Mgat3 gene (**Figure 2G**). Similarly, the high expression of Fut9 in kidney and brain regions also aligned with our observation of elevated fucosylated *N*-glycans in these tissues (**Figure 2H**). Collectively, these findings delineate a detailed and tissue-resolved map of the tissue-specific *N*-glycan patterns, highlighting the complex and diverse glycosylation in mouse tissues.

## 3. Subcellular Localization Constrains the Conservation and Diversification of *N*-Glycosylation

To investigate the underlying determinants of glycosylation variability, we examined whether subcellular localization contributes to the diversities of site-specific glycans. Subcellular localization analyses revealed that glycoproteins identified at the glycopeptide level were predominantly located in the endoplasmic reticulum (444), Golgi apparatus (311), plasma membrane (169), and etc (**Table S5**; **Figure 3A**). Notably, distinct glycosylation patterns were observed across these compartments (**Figure 3B**), highlighting compartment-specific regulation of *N*-glycan biosynthesis^34^. For example, glycoproteins in the endoplasmic reticulum (ER) were mainly decorated with oligo-mannose glycans, characterized by simple branch structures composed solely of hexose (Hex) and unbranched residues, consistent with early-stage glycan biosynthesis^34^. In terms of core structures, core I was most abundant in the ER. In contrast, glycoproteins associated with the extracellular matrix (ECM) and plasma membrane predominantly carried complex glycans featuring extensive branch structures and terminal capping such as sialylation and fucosylation, indicative of advanced processing in the Golgi apparatus (**Figure 3B**). Consistently, core II was the dominant structure in the ECM and plasma membrane compartments. GO enrichment analysis further revealed that ER-localized glycoproteins performed conserved functions, including mediating protein maturation, quality control, and vesicular trafficking, while plasma membrane glycoproteins support more diverse roles regarding signal modulation, ion transport, and cell-type-specific processes—functions that may demand higher structural variability and contextual responsiveness (Figure 3C).

**Figure 3.**
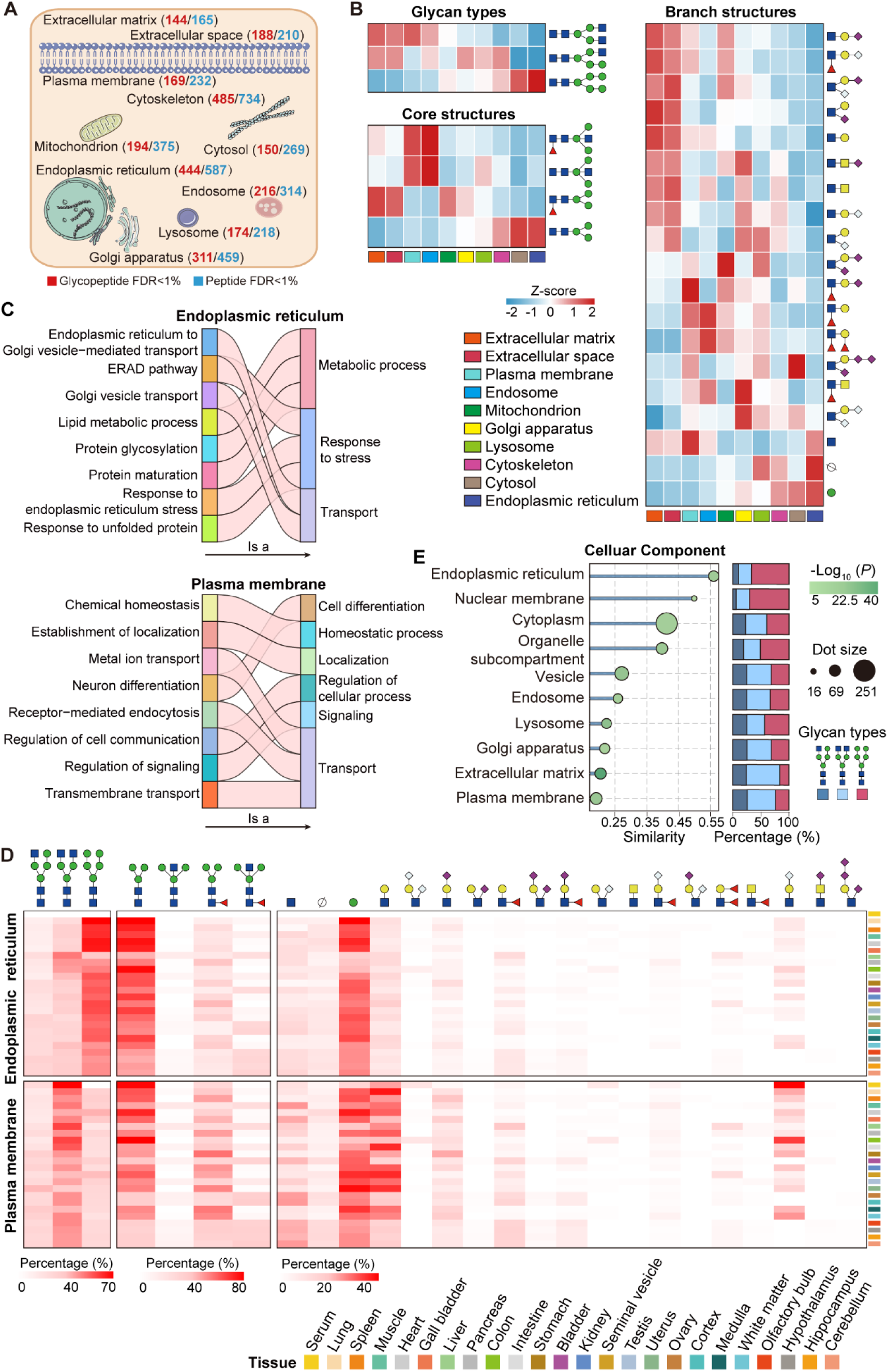
Conservation and Diversities of *N*-Glycans among different subcellular localizations. (A) Subcellular localization of mouse glycoproteins identified at the peptide level or at both peptide and glycan levels (FDR<1%) (see also Table S5). (B) Glycan structural features associated with glycoproteins in distinct subcellular compartments. (C) Gene Ontology (GO) enrichment analyses of glycoproteins from distinct compartments. Cross-tissue comparison of compartment-specific glycosylation patterns within the endoplasmic reticulum (ER) and plasma membrane. (D) *N*-Glycan structural features associated with glycoproteins in ER and plasma membrane across 24 tissues. (E) Glycan similarity analysis of glycoproteins in 12 arbitrary tissues across distinct subcellular compartments.

To examine whether glycosylation conservation or diversities is linked to subcellular compartmentalization across tissues, we specifically compared the glycan profiles of ER- and plasma membrane-localized glycoproteins in multiple tissues and serum. Although tissue-specific differences were still evident, the overall compartment-based glycan feature remained highly conserved. Specifically, *N*-glycan maturation followed a consistent intracellular-to-extracellular trajectory, progressing from oligo-mannose to structurally complex glycans (**Figure 3D**). To quantify the extent of conservation within compartments, we selected shared glycoproteins identified in ≥12 tissues, categorized them by subcellular location, and assessed inter-tissue glycan similarity. ER-localized glycoproteins exhibited the highest glycan similarity across tissues, reflecting conserved biosynthetic constraints. Conversely, ECM and plasma membrane glycoproteins showed significantly greater glycan heterogeneities, consistent with their specialized extracellular functions (**Figure 3E**). Overall, the subcellular localization of glycoproteins shapes *N*-glycosylation patterns, with increasing structural complexity from the ER to the extracellular compartments.

## 4. Diverse Glycosylation Remodels Common Glycoproteins Across Tissues

Building on the observation that *N*-glycosylation patterns vary markedly across tissues, we next examined whether conserved glycoproteins undergo tissue-specific glycan remodeling to support functional specialization. Among the 62 glycoproteins ubiquitously detected, 24 were expressed in all tissues, while other 38 were only absent from serum. Notably, 51 glycoproteins were exclusively detected in neural tissues, suggesting a conserved and specialized glycoproteomic signature within the nervous system (**Figure 4A**). To further explore how a common proteome is customized by glycosylation across tissues, we compared the *N*-glycan profiles of these conserved glycoproteins across at least 12 arbitrary tissues. Despite the invariant protein backbones, the attached *N*-glycan structures varied markedly among tissues, revealing extensive tissue-specific glycan remodeling (**Figure 4B**). This pattern was further validated using an extended set of 24 universally expressed glycoproteins, reaffirming the reproducibility and biological relevance of these tissue-specific glycosylation profiles (**Figure S6**). Remarkably, analyzing glycan variants on 24 shared glycoproteins yielded more robust and specific tissue signatures than global glycan profiling. For instance, a trisialylated *N*-glycan structure was highly abundant in the hippocampus but minimal present in the medulla, olfactory bulb, and hypothalamus, a regional distinction less evident in global glycan profiling, suggesting that differential glycosylation of commonly expressed proteins offers a more sensitive and precise readout of tissue-specific glycan features than total glycan analysis. To further investigate the basis of this remodeling, we categorized the commonly expressed glycoproteins by their subcellular localization and assessed glycan similarity within each category across tissues. Consistent with our earlier findings, glycoproteins localized to the ER exhibited the highest glycan similarity, while those at the cell surface and in the ECM displayed the lowest similarity, reaffirming that glycosylation patterns are shaped by both compartmental constraints and functional demands (**Figure 4C**).

**Figure 4.**
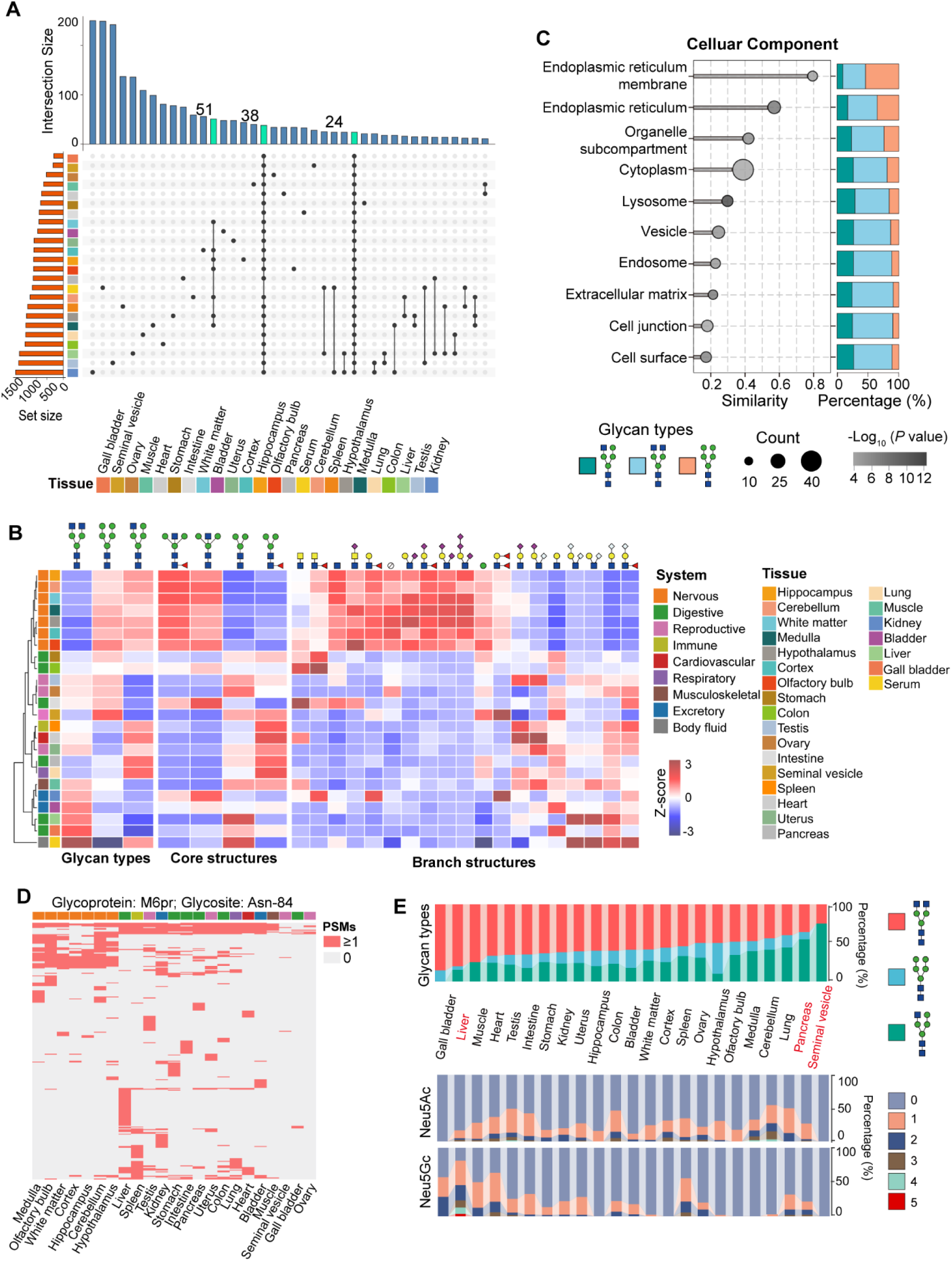
Comparison of glycan structures at commonly expressed glycoproteins across tissues. (A) Upset plots showing shared and unique glycoproteins identified in each tissue. (B) Distribution of different structural modules of site-specific *N*-glycans from glycoproteins identified within at least 12 tissues. The percentages were calculated based on unique glycopeptides containing each type of glycan structural feature versus total glycopeptides. (C) Glycan similarity analysis of widely expressed glycoproteins across distinct subcellular compartments. (D) Tissue-specific glycosylation patterns at the glycosite Asn-84 of mannose-6-phosphate receptor (M6pr), illustrating substantial structural diversity across tissues. (E) Representative glycan types and the number of sialic acid (Neu5Gc and Neu5Ac) residues at the glycosite Asn-84 of M6pr across tissues.

To further explore how *N*-glycosylation modulates protein function in a tissue-dependent manner, we performed site-specific glycosylation analysis on these commonly expressed glycoproteins. The mannose-6-phosphate receptor (M6pr), which exhibited the highest number of *N*-glycans on the common glycosite among the non-serum shared glycoproteins, served as a representative example. Glycosylation at Asn-84 of M6pr displayed remarkable heterogeneities across tissues, with distinct *N*-glycan structures found even at this single glycosite depending on tissue types (**Table S6**; **Figure 4D**). To probe the functional relevance of this site-specific glycan remodeling, we aligned glycan heterogeneities with known tissue-specific roles of the corresponding proteins. Although the amino acid sequences of these glycoproteins remain invariant, variations in their *N*-glycan structures likely modulate their biochemical properties and functional outputs in a tissue-dependent manner. As an example, M6pr is essential for lysosomal enzyme trafficking and is broadly expressed across tissues^35^. However, the glycosylation at the glycosite Asn-84 varied substantially among tissues. In the liver, M6pr predominantly carried complex glycans with extensive sialylation (**Figure 4E**), a modification associated with prolonged protein half-life and enhanced endosomal recycling^36^. In contrast, in seminal vesicle and pancreas, Asn-84 was mainly decorated with hybrid glycans with sialylation (**Figure 4E**). Collectively, tissue-specific remodeling of glycan structures on shared proteins exemplifies how post-translational modifications adapt core biological machinery to diverse physiological demands.

## 5. Site-Specific *N*-Glycosylation display outstanding performance in Tissue Identity

Correlation and unsupervised clustering analyses demonstrated that glycopeptides outperform both glycoproteins and total glycans in distinguishing tissues, underscoring the superior discriminatory power of site-specific glycosylation profiling. Clustering heatmaps and principal component analysis (PCA) plots based on glycopeptide data revealed distinct tissue-specific patterns. Notably, all seven brain regions clustered tightly together, exhibiting glycosylation profiles markedly different from other tissues. Serum formed a distinct cluster, while anatomically adjacent organs grouped more closely, reflecting shared physiological functions. Conversely, organs with distinct roles, such as those within the female reproductive system, displayed greater divergence (**Figures 5A** and **5B**). Functional specialization was further reflected by the abundance of key glycoproteins in specific tissues (**Figure S7**). In the nervous system, the predominant glycoproteins included Slc1a2, Thy1, and Syp, all of which are critical for synaptic transmission, neuronal signaling, and synaptic integrity^37, 38^. In contrast, Muc2 was highly expressed in the intestine and colon, consistent with its role in mucus layer formation, gut homeostasis, and pathogen defense^39^ (**Figure 5C**).

**Figure 5.**
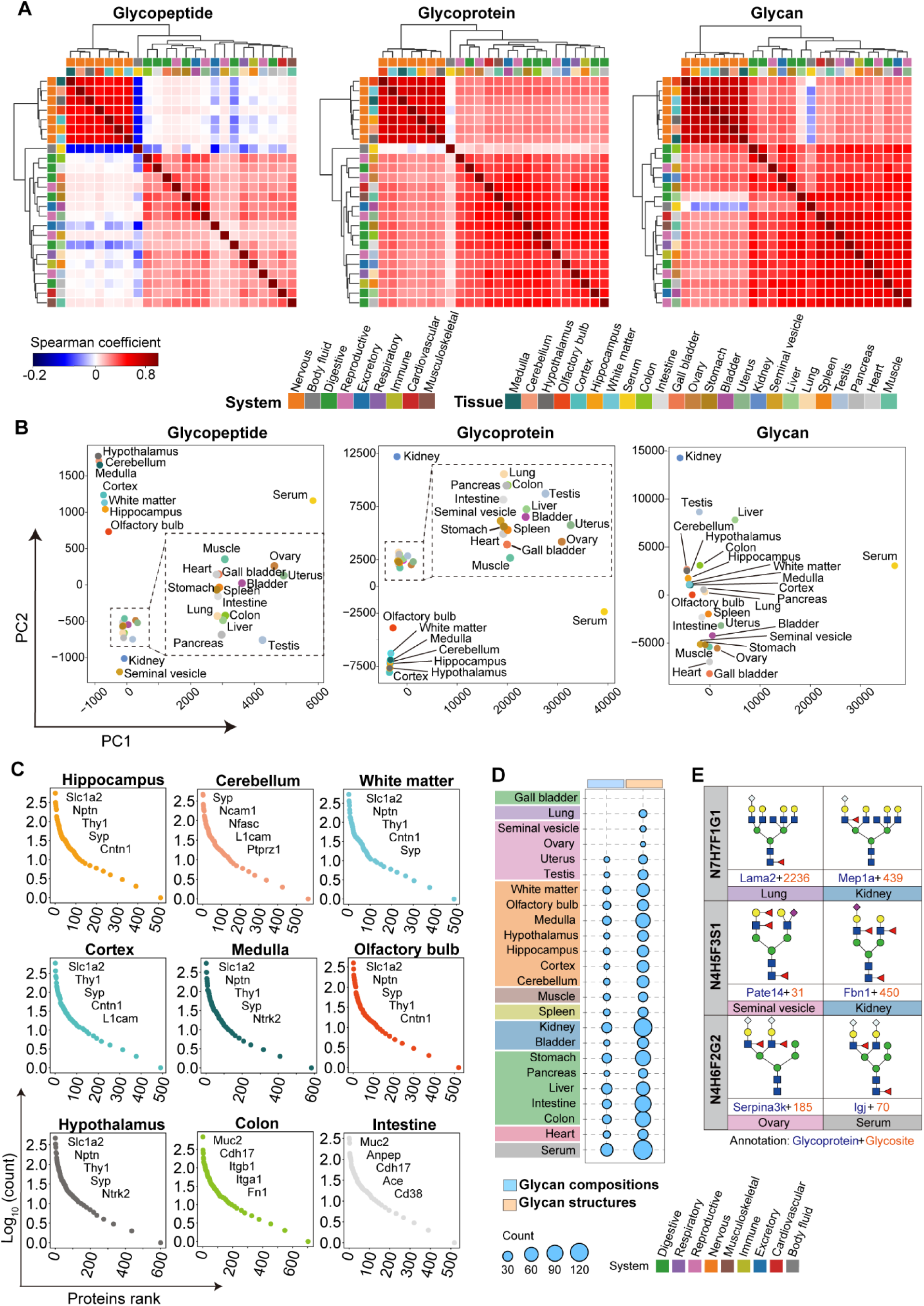
Site-specific *N*-glycosylation as a molecular signature for tissue differentiation. (A and B) Heatmaps and principal component analysis (PCA) plots showing unsupervised hierarchical clustering based on glycopeptide, glycoprotein, and glycan profiles across 24 tissues. (C) Dynamic range of the intensity-ranked glycoproteins within 24 tissues. Five of the high abundant glycoproteins that relate to the functional specialization of the corresponding tissue are listed in descending order. (D) Bubble plots showing the unique glycan composition and structure identified in each tissue. (E) Representative examples of glycan isomers that can distinguish different tissues.

Beyond individual glycoproteins, distinct tissue-specific glycan compositions and structures were observed across 20 and 23 tissues, respectively. These abundant glycan features served as molecular signatures that robustly distinguishing tissues and serum. Brain regions, for instance, exhibited unique fucosylated glycoforms mostly containing multiple Neu5Ac residues, including N5H6F1 in the cerebellum, N4H6F2S3 in the cortex, N4H5F1S6 in the hippocampus, N4H7F3S2 in the hypothalamus, N8H6F1S4 in the medulla, N4H7F3S1 in the olfactory bulb, and N6H7F4S3G1 in the white matter.

In contrast, serum displayed highly specific glycan compositions such as N5H5G4, N5H7G2, and N5H8G1, whereas the gallbladder lacked unique glycan features, likely due to its lower identification coverage (**Figure 5D**). Importantly, glycan structural information provided greater resolution for tissue discrimination than compositional data alone. For example, although the lung, seminal vesicle, and ovary could not be clearly distinguished based on glycan compositions, they exhibited distinct structural isomers, highlighting the importance of glycan isomerism in tissue-specific differentiation (**Table S7; Figure 5E**). These isomeric forms were further confirmed by their characteristic B/Y ions in MS2 spectra (**Figure S8**). Together, these findings highlight the role of site-specific *N*-glycosylation patterns and glycan structural isomers as molecular signatures driving tissue identity.

## 6. Tissue-Enriched Glycans Prefer to Perform Specialized Functions

To investigate the tissue-specific distribution and functional roles of sialylation and fucosylation, which usually play essential role in mammalian development^40^, we systematically quantified the number of fucose and sialic acid residues on *N*-glycans across 24 mouse tissues. Fucosylated *N*-glycans were ubiquitous across all tissues, with particularly high enrichment in the brain regions (average 64.5%), kidney (63%), and seminal vesical (59.1%) (**Figure 6A**). In contrast, sialylated glycans exhibited a more heterogeneous distribution. Although both the kidney and seminal vesicle displayed elevated fucosylation, the seminal vesicle showed relatively low sialylation (12.5%). Notably, the composition of sialic acid varied by tissues, with Neu5Gc demonstrating greater tissue specificity than Neu5Ac. Neu5Gc was virtually absent from neural tissue but abundant in serum and liver (**Table S8**; **Figures 6B** and **6C**). The stomach exhibited the most extensive fucosylation per glycan, with up to nine fucose residues per structure, while the cerebellum, hippocampus, uterus, and liver showed the highest levels of sialylation per glycan, each with distinct Neu5Ac/Neu5Gc profiles (**Figure 6D**). Across all tissues, the average Neu5Gc-to-Neu5Ac ratio was 0.6, but extreme values were observed in serum (18.5), liver (4.3), ovary (3.4), and neural tissues (0.03-0.5) (**Figure 6E**). These observations reflect tissue-specific glycan structures and reaffirm that the murine brain *N*-glycome is dominated by Neu5Ac with only trace Neu5Gc, while serum is rich in Neu5Gc^41^.

**Figure 6.**
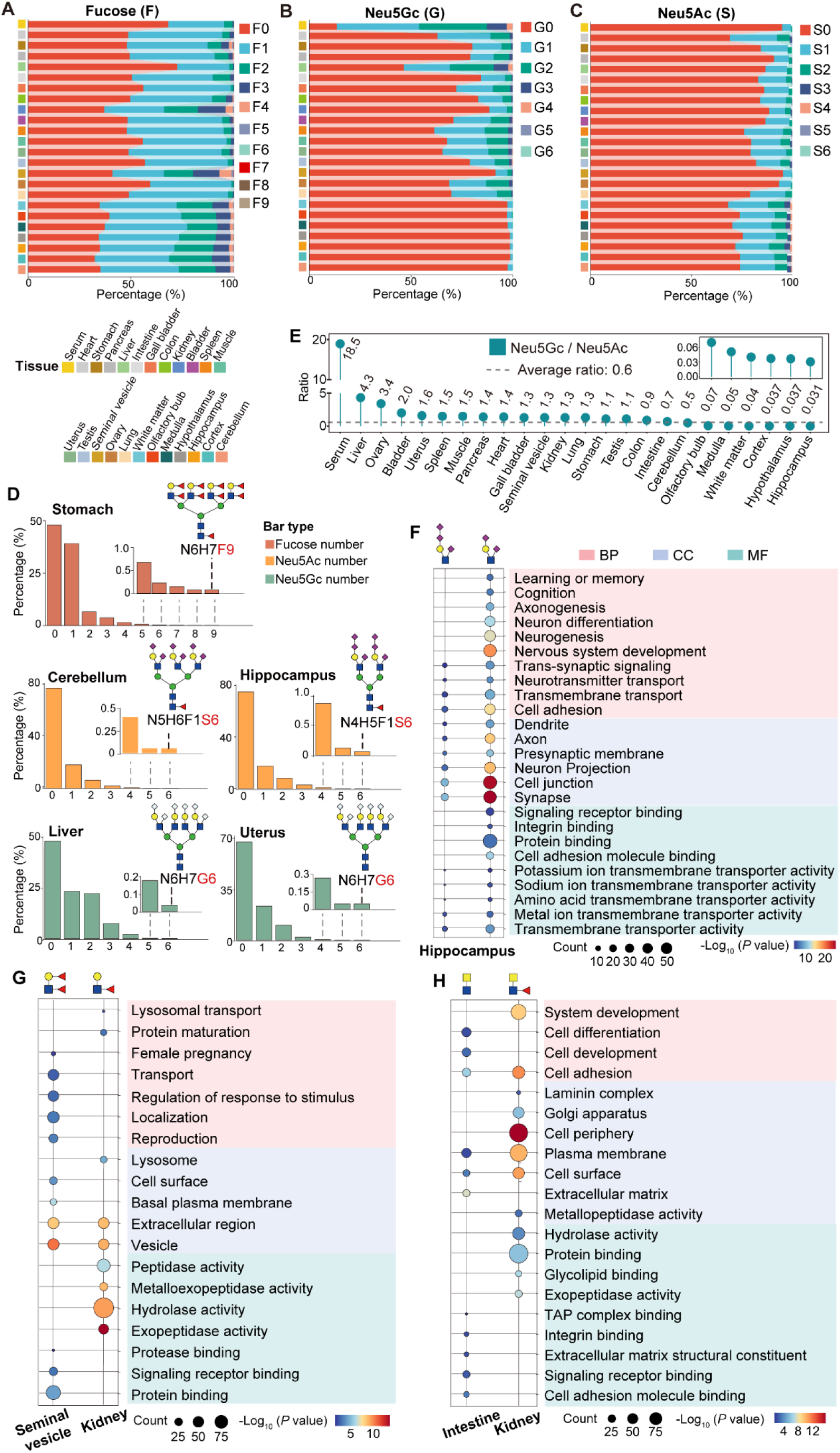
Functional analyses of tissue-enriched glycans across 24 mouse tissues. (A-C) Proportion of different numbers of fucoses and and sialic acid (Neu5Gc and Neu5Ac) residues on intact glycopeptides (IGPs) in each tissue. (D) Representative glycan structures that contain the highest number of fucose or sialic acid residues in five different tissues. (E) The ratio of Neu5Gc-to-Neu5Ac in each tissue. Average ratio among 24 tissues was 0.6. (F-H) Gene ontology enrichment of glycoproteins modified by (F) disialylated and trisialylated structures in the hippocampus, (G) Lewis^x/a^ and Lewis^y/b^ epitopes in the kidney and seminal vesicle, respectively; and (H) LacdiNAc-containing structures in the colon and kidney.

Ploy-sialic acid (ploySia) glycans, particularly those modified by Neu5Ac, were highly enriched in all seven brain regions, with considerable structural variations in branches (**Figure S9A**). The hippocampus showed robust expression of polySia, including disialylated and trisialylated forms. Gene Ontology (GO) analysis indicated that glycoproteins modified by trisialylated glycans were predominantly localized to synapses, presynaptic membrane, and axon, implicating them in cell adhesion, transmembrane transport, and trans-synaptic signaling. Disialylated glycoproteins, also abundant in the hippocampus, were similarly involved in transmembrane transporter and cell adhesion, and were additionally enriched for functions related to cognition, learning, and memory (**Figure 6F**). Interestingly, sialylated LacdiNAc glycans clustered more closely with ploySia structures than with other LacdiNAc glycans categorized by branch features (**Figure 2E**). These sialylated LacdiNAc structures showed highest abundance in the olfactory bulb and were associated with neuron differentiation and synapse organization, with a subcellular localization pattern similar to that of polySia glycans (**Figure S9B**).

Tissue-specific enrichment was also observed for Lewis epitopes. Lewis^x/a^ was abundant in the kidney, while Lewis^y/b^ was preferentially expressed in the seminal vesicle (**Figures 2E and S9C**). GO enrichment analysis revealed that Lewis^x/a^ modified glycoproteins in the kidney localized to vesicles, extracellular region, and lysosome, and were functionally linked to peptidase activity, protein maturation, reproduction, and lysosomal transport. In parallel, Lewis^y/b^ modified glycoproteins in the seminal vesicle shared similar localization patterns and were associated with protein binding and signaling receptor binding, with significant implications for reproduction, response to stimulus, female pregnancy (**Figure 6G**). Notably, sialylated Lewis epitopes with Neu5Ac or Neu5Gc preferentially clustered with glycans carrying the same type of sialic acid (**Figure 2E**).

LacdiNAc glycans were most enriched in the kidney, colon, stomach, and intestine. Among these, the intestine had the highest abundance levels of LacdiNAc structure, whereas the kidney exhibited the most abundant fucosylated LacdiNAc (**Figure S9D**). In the intestine, LacdiNAc-modified glycoproteins were primarily localized to the extracellular matrix, cell surface, and plasma membrane, with functions related to cell adhesion, cell development, and differentiation. In the kidney, fucosylated LacdiNAc glycoproteins were enriched at the cell surface and in the laminin complex, where they were linked to peptidase activities, cell adhesion, and organ development (**Figure 6H**). These results indicate that specialized glycan structures are preferentially attached to functionally relevant glycoproteins in specific tissues, suggesting a coordinated mechanism by which glycan modifications fine-tune protein function to meet localized physiological demands.

## 7. Site-Specific *N*-Glycan Show Non-Random Microheterogeneities

To investigate prevalent patterns of intact *N*-glycans at specific glycosites, we performed a systematic analysis of co-occurring possibility across 24 mouse tissues. Specifically, to ensure data robustness, we used stringent filtering criteria including glycosites with at least 10 distinct glycan structures, with each glycan detected a minimum of 10 times per tissue. Glycan pairs were further filtered by requiring a co-occurrence probability ≥ 0.5 at the same glycosite. Based on these criteria, we systematically examined 24 mouse tissues, focusing on six predominant co-occurrence patterns as well as other combinations, including (i) isomeric glycans; (ii) glycans differing by one or more sialic acid residues; (iii) glycans differing by one or more fucose residues; (iv) glycans differing by one or more mannose/ galactose residues; (v) glycans differing by both fucose and sialic acid residues; and (vi) glycans differing by a mannose or galactose residue with additional random variations in fucose and sialic acid content. Among these patterns, pattern (ii)—glycans differing by sialic acid content—was particularly common in several tissues. In muscle, for example, 63.3% of co-occurred glycan pairs differed in sialic acid residues, so as the gallbladder (53.3%). Pattern (iii), involving differences in one or more fucose residues, was notable in cerebellum (29.1%) and seminal vesicle (26.7%) (**Figure 7A**). These findings suggest that the site-specific glycan microheterogeneities may often arise from dynamic additions or removals of terminal sialic acids and fucose, rather than from divergent biosynthetic pathways.

**Figure 7.**
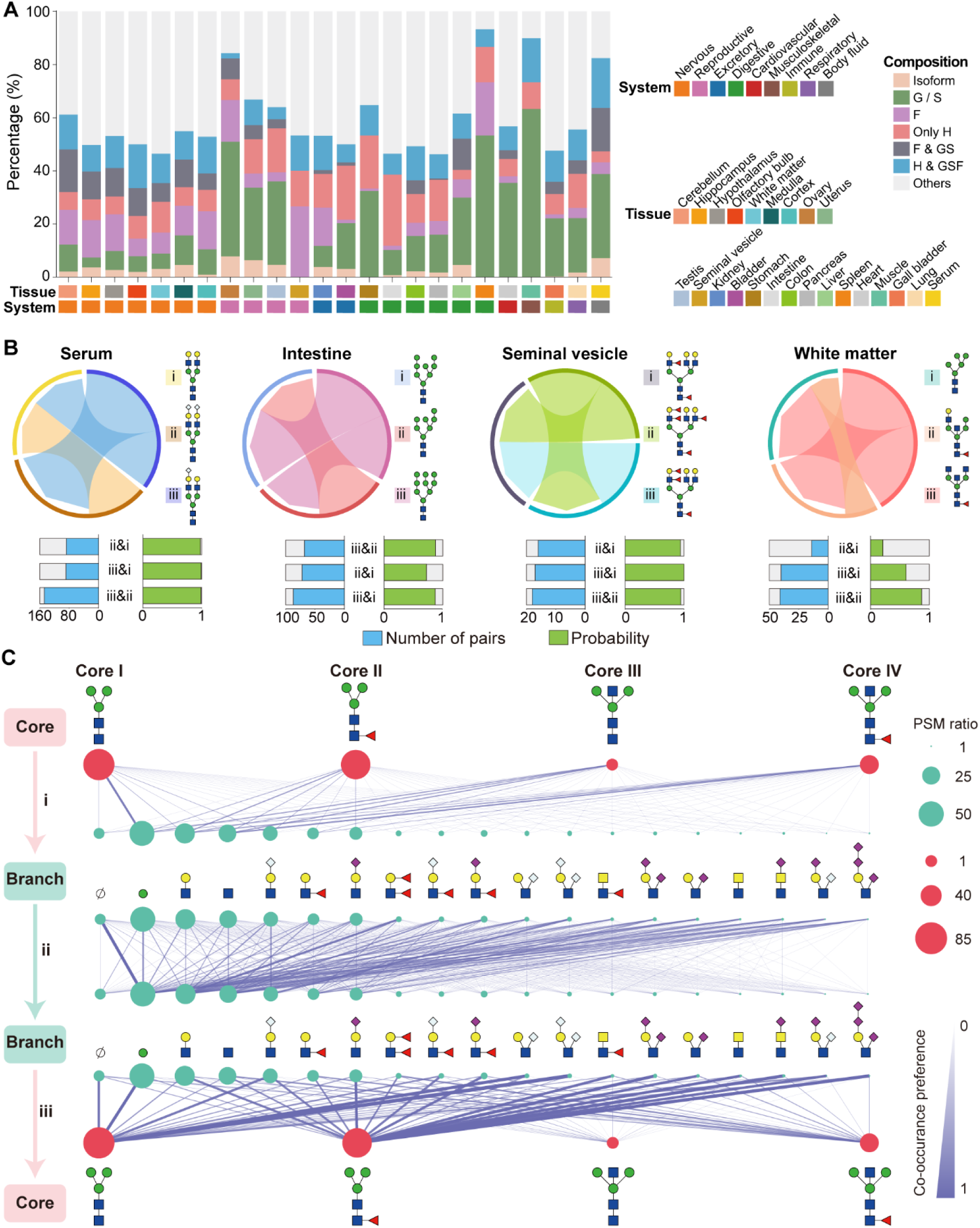
Co-occurrence network analyses on site-specific *N*-glycan microheterogeneities and preference analyses on modular Structural Features of *N*-Glycans. (A) Summary of six predominant glycan co-occurrence patterns across 24 mouse tissues. Bar plots represent the proportion of glycan pairs differing by one or more sialic acid, fucose, or other monosaccharide units. Hexose (H), Fucose (F), Sialic acid (S), and N-acetylglucosamine (G). Isoform: Isomeric glycans. G/S: differ only in G or S. F: differ only in F. Only H: Differ only in H. F & GS: differ in both F and either G or S. H & GSF: Differences in H and at least one of G, S, or F. Others: All other unclassified variations. (B) Examples of co-occurrence patterns and co-existance probabilities of intact *N*-glycans at individual glycosites in representative mouse tissues (see also Figure S10). (C) Associations between core and branch structures at site-specific level across 24 mouse tissues. The network diagrams illustrate co-occurrence probabilities between glycan structural elements based on uniquely identified glycopeptides in the mouse tissues. For each panel: (i) Core-to-branch associations depict the likelihood of specific branch structures occurring alongside distinct core structures. (ii) Branch-to-branch associations show how frequently different branch structures co-occur on the same glycan. (iii) Branch-to-core associations indicate the probability of specific core structures appearing with given branch structures. Edge darkness reflects the strength of co-occurrence (darker lines = higher probability), and node size corresponds to the frequency of each structural element. For branch structures with repeats, each structure is counted only once per glycan (see also Figure S11).

To further illustrate the pattern of dynamic additions or removals of terminal sialic acids and fucose, we examined tissues enriched in sialylated, fucosylated, or mannose glycans. For each tissue, we identified the most abundant glycan structure and calculated its conditional co-occurrence probabilities with others. In serum, the most frequently observed glycan was a biantennary structure carrying two terminal sialic acid residues (HexNAc_4_ Hex_5_Fuc_1_NeuAc_2_, N4H5F1S1). Notably, when this structure was detected at a glycosite, up to 148 other biantennary structures containing one sialic acid was concurrently identified at the same site, with a co-occurrence probability of 98.7%. Additionally, 88 asialylated biantennary structures were also frequently present, with a co-occurrence probability of 97.8%. Furthermore, a total of 89 pairs of monosialylated and asialylated biantennary glycans co-occurred at individual sites with a high probability of 98.9% (**Figure 7B**). To assess the generality and tissue specificity of this co-occurrence pattern, we extended our analysis to the remaining 23 mouse tissues. These triplets (di-, mono-, and asialylated glycans) were reproducibly identified across tissues such as liver, heart, ovary, and muscle (**Figure S10A**). Interestingly, while both serum and liver exhibited this patterns, their dominant glycan structures differed. In serum, disialylated and monosialylated biantennary glycans predominated, whereas asialylated counterparts were more abundant in liver. This divergence may reflect tissue-specific glycoprotein turnover requirements: sialylation is known to prolong the circulatory half-life of glycoproteins^42^, while the absence of terminal sialic acids facilitates rapid clearance and renewal of proteins in the liver, thereby supporting its metabolic functions.

In addition to the sialylated biantennary glycans, similar co-occurrence patterns were also observed for mannosylated and fucosylated glycans in tissue-specific contexts. In both small intestine and testis, numerous glycosites exhibited the simultaneous presence of oligo-mannose differing by only one or a few mannose residues. For example, Man7, Man8, and Man9 frequently co-occurred at the same sites in the small intestine, often appearing in pairs (**Figure S10B**). Likewise, in the seminal vesicle and kidney, fucosylated glycans displayed a similar trend: glycans differing by one or more fucose residues frequently coexisted at individual sites (**Figure S10C**). Notably, a distinct pattern was observed in the mouse brain, where all seven brain regions predominantly expressed bisected complex glycans modified with core fucose (Core IV). However, unlike other tissues where co-occurring glycans typically varied by single monosaccharide units, glycosites in the mouse brain exhibited more heterogeneous and structurally diverse co-occurrence profiles, which further supports our earlier observation that nervous system featured the greatest structural diversity. Specifically, in the white matter, the three most frequently coexisting glycans with the core-fucosylated bisecting glycans represented distinct glycan types—complex, hybrid, and oligo-mannose—each characterized by short and simple configurations (**Figures 7B and S10D**). These results underscore the unique glycosylation landscape of the brain, where co-occurrence patterns reflect a broader range of biosynthetic pathways and structural motifs than those observed in other tissues. Overall, our findings provide a detailed and novel perspective on the structural microheterogeneities of *N*-glycans, offering new insights into the dynamic nature of glycosylation across tissues, which has not been systematically addressed in previous studies.

## 8. Co-occurrence Preferences Across Modular Structural Features of *N*-Glycans

We further explore the interplay between core and branch structure of *N*-glycans through calculating the probability of specific branch structures co-occurring with particular core structures, as well as the likelihood of co-occurrence between different branch structures. Core I showed a strong preference for mannose (Hex), consistent with oligo-mannose characteristics, while Core II exhibits a notable preference for LacNAc (HexNAc+Hex). Core III and Core IV, both of which represent bisected core structure, tend to associate with branch structures lacking sialic acid or fucose capping^30^ (**Figures 7C-i** and **S11A**). Branch-branch associations revealed more intricate patterns. Branches with more complex structures (on the right side of the network) had a higher probability of co-occurring with either a single GlcNAc or LacNAc. Additionally, ploySia-containing branches frequently co-occurred with Lewis^x/a^ structures, while fucosylated branches tended to cluster together. Sialylation on one branch increased the likelihood of sialylation on the opposite branch, with Neu5Ac and Neu5Gc modifications preferentially pairing with their respective types (**Figures 7C-ii** and **S11B**). Further analysis of branch-to-core structure associations uncovered additional insights. Branch types such as the unbranched structure (labeled as Ø), Hex, and Neu5Gc-modified structures show a stronger preference to core I, whereas LacNAc, Neu5Ac-modified branches, and fucosylated branch structures preferentially attach to core II. The low abundance bisected *N*-glycan structures, core III and core IV, exhibited minimal branch association, except with simple structures like GlcNAc. This might be due to the inhibitory effect of bisected glycans on key branching glycosyltransferases (GnTII, GnTIV, and GnTV) and fucosyltransferase (FUT8), limiting *N*-glycan branching and elongation^30^ (**Figures 7C-iii and S11C**). Together, these findings reveal that substructural co-occurrence patterns are shaped not only by terminal modifications of *N*-glycans but also by intrinsic constraints between core and branch elements, contributing to the observed glycan microheterogeneity across tissues.

## Discussion

In this study, we present the first high-resolution atlas of the *N*-glycoproteome across 24 tissues, encompassing 16 major organs, 7 distinct brain regions, and serum. This comprehensive dataset provides an in-depth characterization of tissue-specific *N*-glycan landscapes, offering critical insights into how glycosylation contributes to functional specialization across biological systems. Tissue-enriched glycans are preferentially associated with specialized biological functions, reinforcing the view that glycan modifications fine-tune protein function to meet localized physiological demands. The systematic analysis demonstrated that site-specific *N*-glycosylation served as a molecular signature for tissue identity, demonstrating superior discriminatory power compared to the proteome and glycome. Furthermore, co-occurrence network analyses uncovered non-random microheterogeneities among site-specific *N*-glycans and co-occurrence preferences across structural features within individual glycans. Altogether, this resource significantly expands our understanding of glycosylation dynamics across multiple tissues and lays the foundation for future translational research in biomarker discovery, disease modeling, and therapeutic development.

Through an extensive site-specific glycoproteomic strategy combined with StrucGP software, we identified 74,277 unique intact *N*-glycopeptides (IGPs) occupying 8,681 glycosites, significantly expanding known mouse *N*-glycoproteome. This unprecedented dataset reveals pronounced tissue-specific glycosylation patterns that reflect the functional and biological specialization of diverse organs. Importantly, site-specific *N*-glycosylation outperformed both proteomes^27^ and glycomes^41, 43^ in distinguishing tissue identity, demonstrating greater discriminatory power and robustness to sampling depth. This was particularly evident in the clear segregation of serum and the cohesive clustering of brain regions, underscoring the sensitivity of site-specific glycopeptide features in capturing tissue-specific signatures.

Across tissues, distinct *N*-glycan structural profiles were observed. The nervous system exhibited the highest structural diversity despite lower overall glycan complexity^44^, reflecting the unique demands of neural tissues where specialized glycans likely play critical roles in neural signaling and synapse maintenance^41, 45^. Notably, core structures further showed striking tissue specificity, with core I structures dominating in circulation and metabolic tissues, whereas bisected cores (core III and core IV) were enriched in neural tissues, suggesting specialized regulatory functions in neuronal signaling^46^. The tissue-specific distribution of branch structures, including polySia glycans, Lewis epitopes, and LacdiNAc structures, implies their specialized roles in organ-specific physiological functions. PolySia glycans were predominantly found in the hippocampus and olfactory bulb, consistent with their pivotal roles in neurodevelopmental processes such as synaptic organization, learning, and memory^47^. Lewis^x/a^ and Lewis^y/b^ epitopes were enriched in the kidney^48^ and seminal vesicle^49^, respectively, implicating them in lysosomal transport and reproductive processes. LacdiNAc glycans were preferentially expressed in the kidney and digestive organs, where they may contribute to cell adhesion, microbial defense^50^ and mucosal protection^51^. These structural patterns were closely linked to tissue-specific glycosyltransferase (GT) expression profiles^52^. For instance, elevated expression of Mgat3 in neural tissues correlated with the abundance of bisected glycans^53^, while fucosylated glycans enrichment in the kidney aligned with high Fut9 expression^54^. These findings provide a molecular basis for the observed tissue-specific glycosylation patterns and emphasize the regulatory influence of GT expression in shaping the glycome.

We further observed that glycosylation profiles were influenced by subcellular localization^55^. Glycoproteins localized to the endoplasmic reticulum display highly conserved glycan profiles, whereas those in the extracellular matrix or plasma membrane exhibited greater variability, transitioning from oligo-mannose to more complex types. This gradient of glycan diversity aligns with the known biosynthetic pathway of *N*-glycans and suggests that site-specific glycosylation adapts to extracellular functional demands^56^. Interestingly, even the same glycosite on a shared glycoprotein exhibited distinct glycan structures across tissues, providing direct evidence that glycoprotein scaffolds can be fine-tuned via tissue-specific glycoforms to meet local functional requirements. Furthermore, structural isomers of the same glycan compositions were often observed in different tissues, which effectively distinguish tissues that cannot be resolved by glycan composition alone, highlighting the added resolution of isomer-level analysis in tissue discrimination.

Co-occurrence network analyses reveal that site-specific *N*-glycan microheterogeneities are governed by non-random, modular co-occurrence patterns shaped by both dynamic terminal editing and substructural preferences. Most co-occurring glycans at the same glycosite differed by terminal residues—particularly sialic acids and fucose—suggesting that local glycan diversity often arises from sequential addition or removal of these monosaccharides rather than divergent biosynthetic pathways. This pattern, especially involving sialic acid variations, was prominent in tissues such as serum, liver, heart, ovary and muscle. where disialylated, monosialylated, and asialylated biantennary structures frequently coexisted with high probability. Further analysis of substructural co-occurrence revealed preferential pairings between specific core and branch structures: core I preferentially associated with mannose branches, core II favored LacNAc and fucosylated branches, while bisected cores (core III and IV) were associated with minimal branching. This corresponds with their known inhibitory effects on key branching glycosyltransferases (GnTII, GnTIV, and GnTV) and fucosyltransferase (FUT8)^57^, indicating structural constraints that limit glycan elongation. Moreover, branch-branch co-occurrence patterns revealed a coordinated regulation of glycan elaboration, with complex branches tended to associate with simpler structures such as single GlcNAc or LacNAc. The frequent co-occurrence of polySia-containing branches with Lewis^x/a^ structures highlights a potential functional synergy, as both modifications are known to mediate cell signaling^58^ and immune evasion^59^.

Despite the breadth of our dataset, several limitations should be acknowledged. The pooling strategy, while necessary for comprehensive organ coverage, may obscure individual variations and low-abundance glycoproteins. Additionally, while the data provide rich glycosylation maps, functional characterization of many identified glycans remains to be explored. Future efforts employing single-cell or spatial glycoproteomics will be essential to resolve cellular heterogeneity and tissue microenvironments. Moreover, expanding this work to disease states, such as aging or cancer, will further illuminate glycosylation dynamics under pathological conditions. Despite of these limitations, the study provides a foundational resource for understanding tissue-specific glycosylation and its broader implications for biomarker discovery, disease modeling, and therapeutic development.

## Methods

### Tissue Preparation

Ten adult C57BL/6N mice (8-9 weeks old; five females and five males) were obtained from Huachuang Xinnuo Pharmaceutical Technology Co. (Jiangsu, China). The mice were anesthetized using avertin and sacrificed following perfusion with 0.9% NaCl. A total 24 tissues, including thw white matter, hippocampus, olfactory bulb, medulla oblongata, hypothalamus, cerebellum, cortex, seminal vesicle, kidney, colon, intestine, testis, bladder, lung, ovary, uterus, pancreas, heart, liver, spleen, stomach, muscle, gallbladder, and serum, were collected for glycoproteomics analysis. All tissues were flash-frozen in liquid nitrogen and stored at -80 °C until protein extraction. All procedures were in compliance with ethical regulations and approved by the ethics committee of Northwest University, China (NWU-AWC-20241206M).

### Protein Extraction and Trypsin Digestion

Each tissue was cut into around 1 mm^2^ fragments and washed three times with precooled phosphate-buffered saline (PBS, pH 7.4). The fragments were then homogenized in 0.5 ml of 8 M urea (cat. no. U5128, Sigma, Germany)/1 M NH_4_HCO_3_ (cat. no. 11213, Sigma, Germany) at 4 °C using a high-throughput tissue grinding machine (Shanghai Jingxin, China). The crude extract was briefly sonicated until the solution was clear. Lysates was clarified by centrifugation at 15,000 g for 15 min at room temperature (RT). Protein concentrations were determined by BCA protein assay reagent (cat. no. P0012, Beyotime, China). All proteins from mouse tissue were reduced by 5 mM dithiothreitol (DTT) (cat. no. 43819, Sigma, Germany) at 37 °C for 1 h, and alkylated by 15 mM iodoacetamide (IAM) (cat. no. I6125, Sigma, Germany) at RT in the dark for 30 min. Another 2.5 mM DTT was then added and incubated at RT for 10 min. The solutions were diluted two times with deionized water prior to the first cycle of protein digestion by incubating the samples with the sequencing grade trypsin (cat. no. V5113, Promega, USA, WI; proteins: enzyme, 100:1, w/w) at 37 °C for 2 h with shaking. The solutions were further diluted four-fold with deionized water and the same relative amount of trypsin (proteins: enzyme, 100:1, w/w) was then added and incubated at 37 °C overnight with shaking. The pH of the solution was adjusted with 10% trifluoroacetic acid (TFA) (cat. no. 3020031, Sigma, Germany) till pH < 2. The samples were centrifuged at 15,000 g for 10 min to remove any particulate matter and the peptides in supernatants were desalted by the C18 column (cat. no. WAT054955, Waters, USA). Peptides were eluted from C18 column in 50% acetonitrile (ACN) (cat. no. A9554, Thermo Fisher Scientific, USA)/0.1% TFA solution and the peptide concentrations were measured by the UV absorbance at 215 nm using a DS-11 spectrophotometer (DeNovix, USA).

### Intact glycopeptide enrichment

The intact glycopeptides were enriched from 24 tissues using two enrichment methods: Oasis Mixed Anion-Exchange (MAX) SPE column^60^ (cat. no. 186000366, Waters, USA), and in-house HILIC micro-column (HILIC beads spec: 5 μm, 100 Å, cat. no. P19-00385, Agela Technologies, China). Briefly, the tryptic peptides from mouse tissue were purified by C18 and were enriched in 50% ACN/0.1%TFA. The peptide solution (∼6 mg peptides) was diluted by 100% ACN and 50% TFA to a final solvent composition of 80% ACN/0.2% TFA. The HILIC micro-column was washed three each by 0.1% TFA and 80% ACN/0.2% TFA, peptide sample and beads were mixed and incubated at room temperature for 2 h, followed by three washes with 80% ACN/0.2% TFA. The glycopeptides bound to the beads were eluted in 0.4 mL of 0.1% TFA solution. About 20 μg glycopeptides were used for LC-MS/MS analysis, and the remaining glycopeptides were used for high-pH HPLC fractionation. The flow through of the HILIC micro-column was loaded onto an anion exchange reversed phase (MAX) column to maximize the glycopeptide enrichment. Briefly, the flow through from HILIC micro-column were diluted by 100% ACN and 50%TFA to the final concentration of 95% ACN/1% TFA. The MAX column was washed three times each by 100% ACN, 100 mM triethylammonium acetate buffer, deionized water, and 95% ACN/1% TFA, followed by sample loading twice and washes four times with 95% ACN/ 1% TFA. The glycopeptides bounding to the column were eluted in 400 μL of 50% ACN / 0.1% formic acid (FA). The sample was dried by RVC 2-18 CDplus concentrator (Christ, Osterode am Harz, Germany) and resuspended in 50 μL of 0.1% FA.

### High-pH HPLC fractionation

HILIC enriched intact glycopeptides from 22 mouse tissues were separately injected via a 1260 Series HPLC (Agilent Technologies, USA) into a Zorbax Extend-C18 analytical column containing 1.8 μm particles at a flow rate of 0.2 mL/min. The mobile-phase A consisted of 10 mM ammonium formate (pH 10) and B composed of 10 mM ammonium formate and 90% ACN (pH 10). Sample separation was accomplished using the following linear gradient: 0-2% B for 10 min, 2-8% B for 5 min, 8-35% B for 85 min, 35-95% B for 5 min, 95-95% B for 15 min. Peptides were detected at 215 nm and 96 fractions were collected along with the LC separation in a time-based mode from 16 to 112 min. The 96 fractions of one tissue were concatenated into six fractions. The samples were then dried in a Speed-Vacuum and stored at -80°C until LC-MS/MS analysis.

### LC-MS/MS analysis

The enriched glycopeptides were separated on an Orbitrap Fusion Lumos Mass Spectrometer equipped with an Easy-nLC^TM^ 1200 system (Thermo Scientific, San Jose, CA, USA). Peptide separation was performed on a 75 μm × 50 cm Acclaim PepMap 100 C18 column (cat. no. 164570, Thermo Fisher Scientific, USA) with a 75 μm × 2 cm pre-column (cat. no. 164946, Thermo Fisher Scientific, USA). The flow rate was kept at 200 nL/min using mobile phase A (0.1% FA in water) and mobile phase B (0.1% FA in 80% ACN). The gradient profile (90 min) was set as follows: 3–7% B for 1 min, 7–25% B for 104 min, 25–35% B for 50 min, 35–70% B for 14 min, 70–100% B for 1 min, 100% B for 10 min. The spray voltage was set at 2.4 kV. Orbitrap MS1 spectra (AGC 4.0×10^5^) were collected from 375 - 2,000 m/z at a resolution of 60 K followed by oxonium ions triggered data-dependent higher-energy collision dissociation (HCD) MS/MS at a resolution of 15 K using an isolation width of 0.7 m/z for 20% collision energy and 2 m/z for 30% collision energy. Charge state screening was enabled to reject unassigned and singly charged ions. A dynamic exclusion time of 20 s was used to discriminate against previously selected ions.

### Intact glycopeptides identification by StrucGP

Intact glycopeptides were identified using the StrucGP software^21^. Briefly, the intact glycopeptide analyses were performed using the built-in glycan branch structure database (containing 17 branch structures) from StrucGP and the *Mus musculus* UniProt protein database (UP000000589, downloaded on 18 Oct. 2023) for the analysis of 24 tissues. The protein enzymatic digestion was set as trypsin with max 2 missing cleavage sites and the potential glycosite-containing peptides were screened with the N-X-S/T motif (X is any amino acid except Proline). Mass tolerances were set to 10 ppm for precursor ions and 20 ppm for fragment ions. For the Y ions determination, an optional mass shift of ±1 Da or ±2 Da was allowed in addition to the 20 ppm mass tolerance in MS2. Finally, 1% false discovery rate (FDR) was set for both peptides and glycans was applied to the quality control of the intact glycopeptide identifications.

### Data quality control via glycan-glycosite correlation analysis

To ensure the reliability and consistency of the glycoproteomic data across all 24 mouse tissues, we performed an internal quality assessment by evaluating the correlation of glycan structures and glycosylation site profiles between technical replicates. Specifically, for each tissue, we selected two representative data frames, either from the same fraction or from max, and calculated Spearman correlation coefficients for (i) the presence of unique glycan structures and (ii) the distribution of glycosylation sites. This correlation-based filtering enabled us to confirm the reproducibility of glycan and site-level profiling within each tissue sample.

### Glycan substructural co-occurrence probability and structural network construction

To investigate structural preferences and co-occurrence patterns among core structures and branch structures, we constructed a substructural-level interaction analysis. First, all unique glycan structures identified across the 24 tissues were decomposed into their constituent motifs. For each glycan, we enumerated motif pairs of three specific types: core–branch, branch–branch, and branch–core. We then quantified the frequency of each motif pair across all glycans and tissues, yielding a global co-occurrence probability matrix. The resulting matrix was visualized as a structural interaction network using NetworkX^61, 62^, enabling intuitive interpretation of motif pairing tendencies and connectivity.

### Glycan-glycan co-occurrence and transition pattern analysis at the glycosite level

At the glycosite level, we analyzed the co-occurrence of glycan structures to assess potential structural transitions and substitution patterns. For this, we restricted the analysis to unique glycosylation sites with a minimum of 10 glycan observations to ensure statistical robustness. For each qualifying glycosite, we counted the number of times each pair of glycan structures co-occurred, resulting in a glycan co-occurrence matrix. A co-occurrence probability matrix was then derived by normalizing the pairwise frequencies against the total occurrence count of the reference glycan within the same site, thereby accounting for asymmetric occurrence likelihoods.

To further dissect potential structural transitions, glycan pairs that satisfied both a minimum co-occurrence count of ≥11 and a co-occurrence probability ≥0.5 were selected for comparative classification. Each glycan pair was categorized into one of seven transition types based on their compositional differences in Hexose (H), Fucose (F), Sialic acid (S), and N-acetylglucosamine (G): (i) Isoform: Isomeric glycans; (ii) G/S: Identical H and F counts; differ only in G or S; (iii) F: Identical H, G, and S counts; differ only in F; (iv) Only H: Differ only in H; identical G, S, and F counts; (v) F & GS: Identical H; differ in both F and either G or S; (vi) H & GSF: Differences in H and at least one of G, S, or F; (vii) Others: All other unclassified variations. This detailed classification system provided insight into the potential biosynthetic flexibility and microheterogeneity at the glycosite level.

### Other bioinformatic analysis

Gene ontology (GO) enrichment analysis was conducted with the Database for Annotation, Visualization, and Integrated Discovery (DAVID) (david.ncifcrf.gov) to identify enriched biological process (BP), cellular component (CC), and molecular function (MF)^63^. Motif was plotted by bioinformatics.com.cn (last accessed on June 20th, 2024), an online platform for data analysis and visualization^64^. Heatmap generation was performed using the seaborn library^65^ from Python, where hierarchical clustering was applied to both rows and columns to reveal global similarity patterns. The linkage method used for dendrogram construction was Ward’s minimum variance method^66^, which minimizes the total within-cluster variance. Distance metrics were based on Euclidean distance. For dimensionality reduction, we applied Principal Component Analysis (PCA) implemented in the sklearn.decomposition^67^ module. All scatter plots, including PCA projections and correlation plots, were generated using matplotlib^68^. Upset plots were performed using UpSetR R package (v 1.4).

## Data availability

The mass spectrometry proteomics data have been deposited to the ProteomeXchange Consortium (proteomexchange.org) via the iProX partner repositor^69^ with the dataset identifier PXD063701. Publicly available RNA-seq data from ArrayExpress (E-MTAB-10276)^4^ were used to assess the expression levels of glycosyltransferases across tissues.

## Code availability

StrucGP software^21^ (v1.2.0) for MS and MS/MS data preprocessing can be downloaded at the Zenodo repository (https://zenodo.org/records/7405508).

## Author contributions

L.J. and S.S. designed the experiments; L.J. performed sample preparation, L.J. performed MS analysis with help from J.L.; Y.W. M.Y., and Y.X., analyzed data and drew figures with the help from X.W., J. Z., Y.C., and Y.Z.; Y.W. and S.S. wrote and edited the manuscript with helps from all authors.

## Acknowledgments

This work was supported by the National Natural Science Foundation of China (Grant No. 22374117), National Key Research and Development Program of China (No. 2019YFA0905200), Natural Science Foundation of Shaanxi Province (Grant No. 2024JC-TBZC-06 and 2023-JC-ZD-46), and Shaanxi Fundamental Science Research Project for Chemistry and Biology (Grant No. 23JHZ006).

