## Supporting information for "A High-Resolution *N*-Glycoproteome Atlas Reveals Tissue-Specific Glycan Remodeling but Non-random Structural Microheterogeneities"

### Supplementary Information

#### 1. Supplementary Figures

**Figure S1** Comparison of glycan compositions identified from 24 mouse tissues in this study with existing glycan databases

**Figure S2** Reproducibility of glycoproteomic data.

**Figure S3** Summary of glycoproteomic identification across 24 mouse tissues.

**Figure S4** LC chromatograph and MS/MS spectra of glycan isoforms identified by StrucGP.

**Figure S5** Representative MS/MS spectra of site-specific *N*-glycans with different antennas.

**Figure S6** Tissue-specific glycosylation of commonly expressed glycoproteins.

**Figure S7** Comparative clustering of glycopeptides, glycoproteins, and glycans across tissues.

**Figure S8** Representative MS/MS spectra of different glycan isoforms identified in different tissues.

**Figure S9** Structural diversity and tissue-specific distribution of functionally enriched glycan epitopes.

**Figure S10** Representative examples of glycan co-occurrence patterns at single glycosites.

**Figure S11** Heatmap showing the co-occurrence preferences across modular structural features of *N*-glycans

#### 2. Supplementary Tables (separate excel files)

**Table S1.** Summary of identified intact *N*-glycopeptides, glycosites, glycans, and glycoproteins across 24 mouse tissues.

**Table S2.** Comparison of identified glycan compositions from 24 mouse tissues in this study with the glycan database-GlyConnect and the glycan databases used by other intact glycopeptide software tools.

**Table S3.** Summary of 3 glycan subtypes, 4 core structures, and 17 branch structures across 24 mouse tissues based on structural modules.

**Table S4.** Expression profiles of 209 glycosyltransferase genes from published RNA-seq data.

**Table S5.** List of glycoproteins enriched in different subcellular localizations in all different mouse tissues.

**Table S6.** Site-specific *N*-glycan heterogeneities at the glycosite Asn-84 of mannose-6-phosphate receptor (M6pr) across mouse tissues.

**Table S7.** Tissue-specific glycan compositions (a) or glycan structures (b) identified across 24 mouse tissues.

**Table S8.** Summary of the number of fucose and sialic acid residues within a single glycan across 24 mouse tissues.

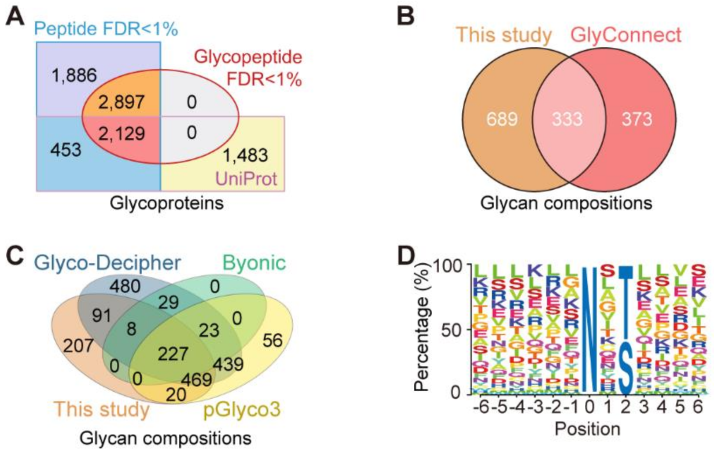


**Figure S1. Comparison of glycan compositions identified from 24 mouse tissues in this study with existing glycan databases.** (A) Comparison of glycoproteins identified from 24 mouse tissues in this study with UniProtKB annotations. The glycoproteins were identified based on their glycosite-containing peptides with FDR<1% at the peptide level or both peptide and glycan (glycopeptide) levels. (B) Comparison of N-linked glycans identified from 24 mouse tissues in this study with the GlyConnect database. (C) Comparison of N-linked glycans identified from 24 mouse tissues in this study with the glycan databases used by the GlycoDecipher, Byonic, and pGlyco3 software (see Table S2). (D) Motif analysis of *N*-glycosites based on FDR<1% at both peptide and glycan levels.

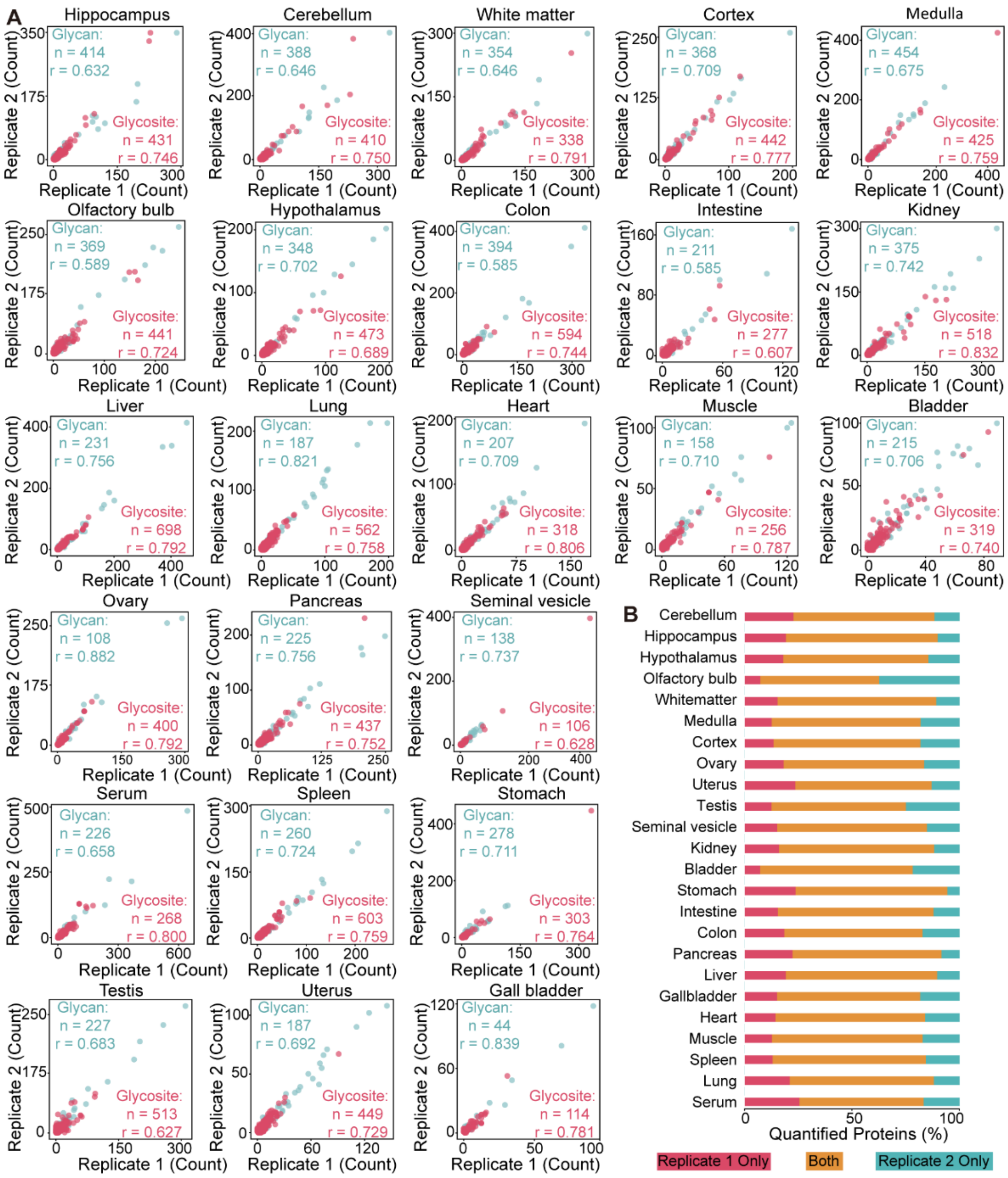


**Figure S2. Reproducibility of glycoproteomic data.** (A) Scatter plots comparing the glycan structure and glycosite identifications between duplicate LC-MS runs across 24 mouse tissues. (B) Overlapped and unique glycoproteins identified by duplicate mass spectrometry analyses of the same glycopeptide sample.

A

| | Glycopeptide Enrichment | | | | |
|---|---|---|---|---|---|
| Tissue | MAX | HILIC | Fraction | PSMs | Unique IGPs |
| Bladder | 3 | 3 | 13 | 36,365 | 6,006 |
| Cerebellum | 3 | 4 | 12 | 65,481 | 8,469 |
| Colon | 3 | 5 | 12 | 59,739 | 10,146 |
| Cortex | 3 | 3 | 12 | 63,212 | 7,803 |
| Gall bladder | 2 | 3 | 0 | 6,375 | 1,304 |
| Heart | 1 | 0 | 12 | 16,271 | 4,642 |
| Hippocampus | 3 | 4 | 12 | 63,733 | 7,623 |
| Hypothalamus | 3 | 3 | 13 | 67,233 | 8,523 |
| Intestine | 3 | 3 | 12 | 33,411 | 6,206 |
| Kidney | 10 | 10 | 15 | 127,201 | 12,208 |
| Liver | 3 | 3 | 12 | 70,026 | 10,862 |
| Lung | 5 | 3 | 12 | 58,255 | 8,139 |
| Medulla | 3 | 3 | 12 | 58,472 | 7,954 |
| Muscle | 3 | 2 | 13 | 19,835 | 3,909 |
| Olfactory bulb | 3 | 3 | 12 | 46,985 | 6,575 |
| Ovary | 6 | 5 | 0 | 22,140 | 2,264 |
| Pancreas | 3 | 4 | 12 | 53,027 | 7,082 |
| Seminal vesicle | 3 | 3 | 12 | 24,742 | 1,660 |
| Serum | 3 | 5 | 18 | 124,034 | 6,903 |
| Spleen | 3 | 3 | 12 | 38,013 | 8,246 |
| Stomach | 4 | 5 | 12 | 22,774 | 4,980 |
| Testis | 5 | 7 | 12 | 79,149 | 8,392 |
| Uterus | 3 | 5 | 16 | 41,174 | 6,581 |
| White matter | 3 | 3 | 12 | 60,890 | 7,531 |

B

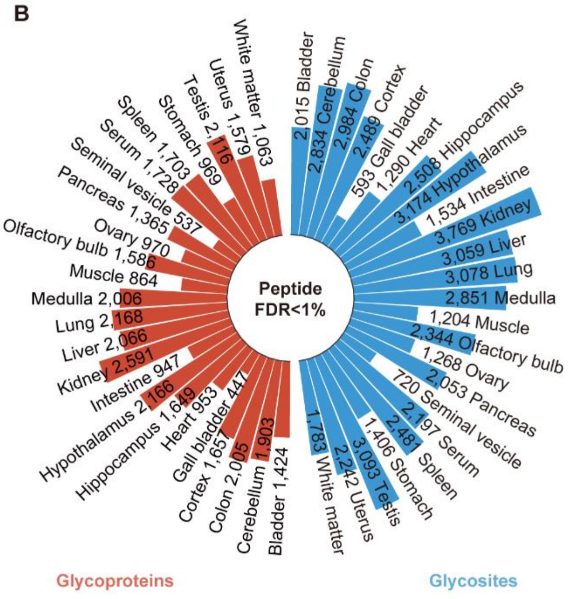


**Figure S3. Summary of glycoproteomic identification across 24 mouse tissues.** (A) Number of identified PSMs and unique IGPs per tissue based on FDR<1% at both peptide and glycopeptide levels. (B) Number of glycoproteins from 24 mouse tissues based on FDR<1% at the peptide level.

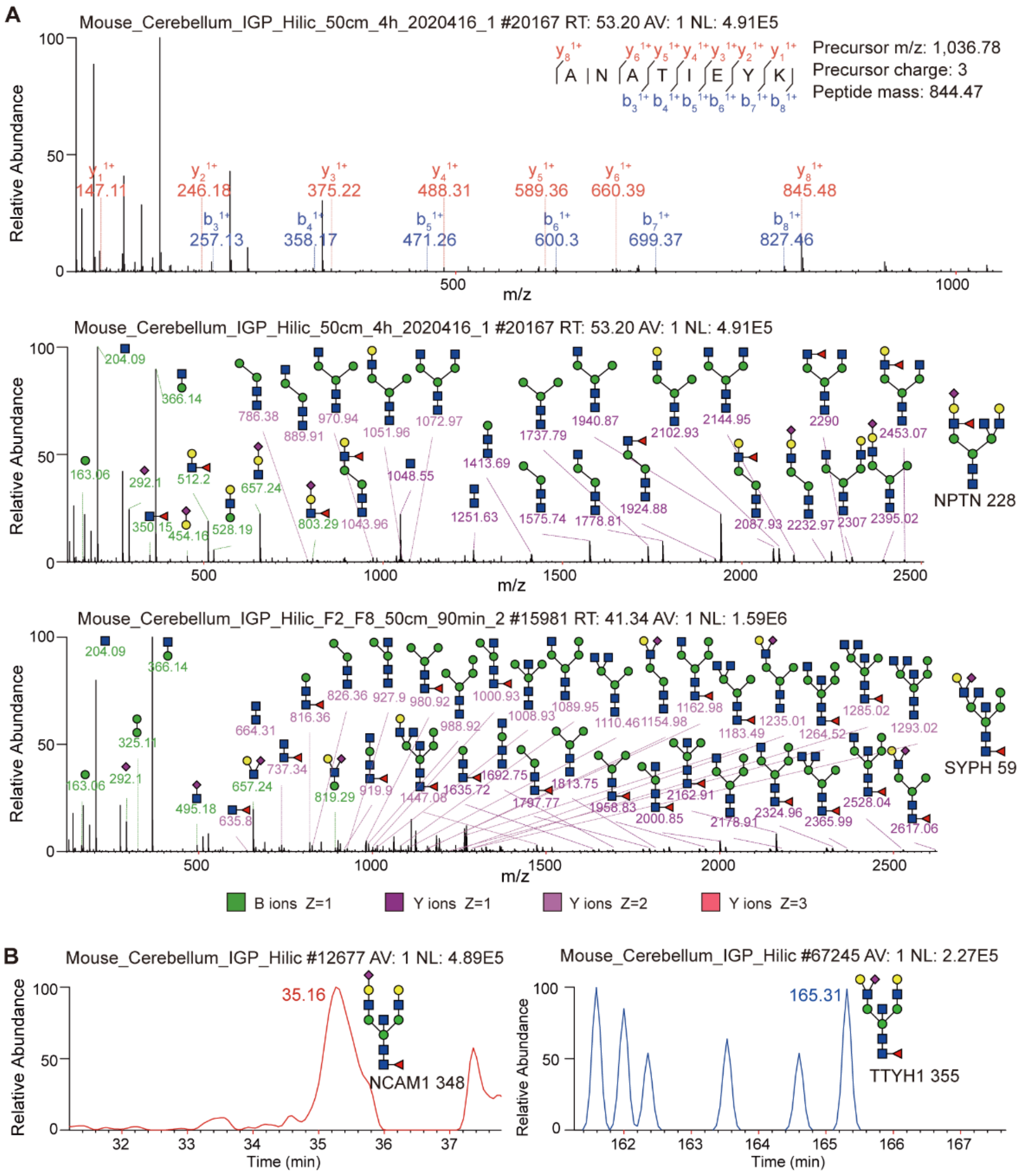


**Figure S4. LC chromatograph and MS/MS spectra of glycan isoforms identified by StrucGP.** (A) Representative MS/MS spectra of 12 structural isomers with the same composition (N5H5F1S1), with only 2 examples shown here, distinguished by characteristic B and Y fragment ions. (B) Chromatographic separation of selected N5H5F1S1 isomers by C18 chromatograph.

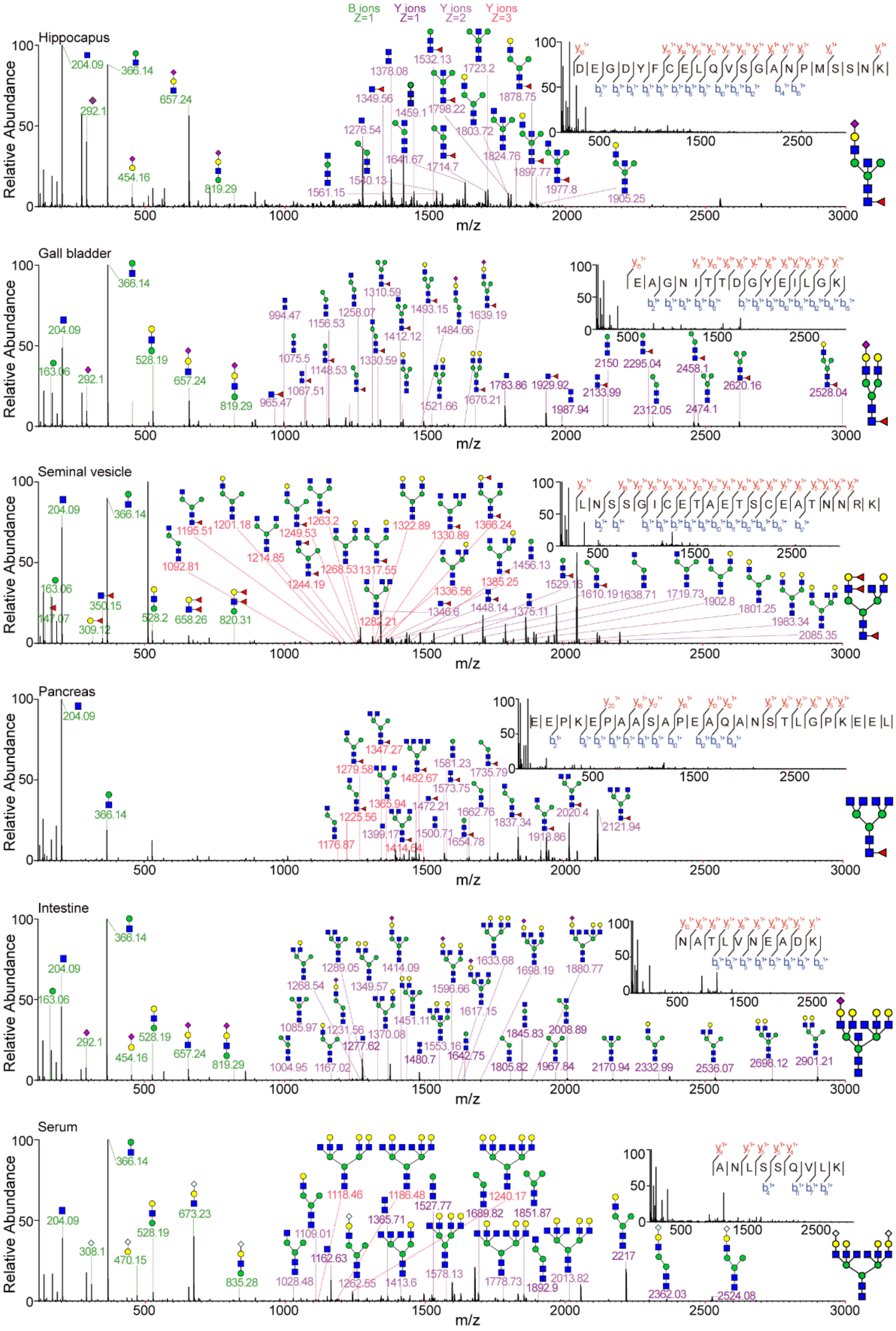


**Figure S5. Representative MS/MS spectra of site-specific *N*-glycans with different antennas.**

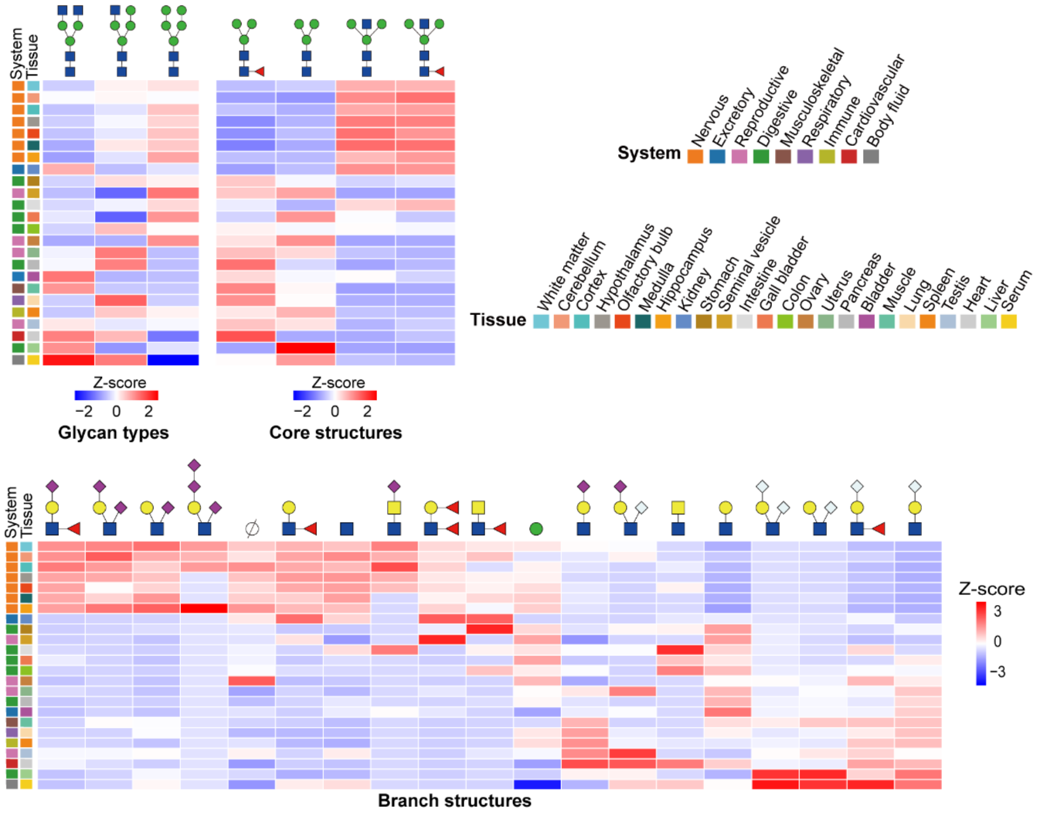


**Figure S6. Tissue-specific glycosylation of commonly expressed glycoproteins.** Heatmap showing site-specific *N*-glycan profiles of 24 glycoproteins commonly expressed in all 24 tissues confirm tissue-dependent structural differences.

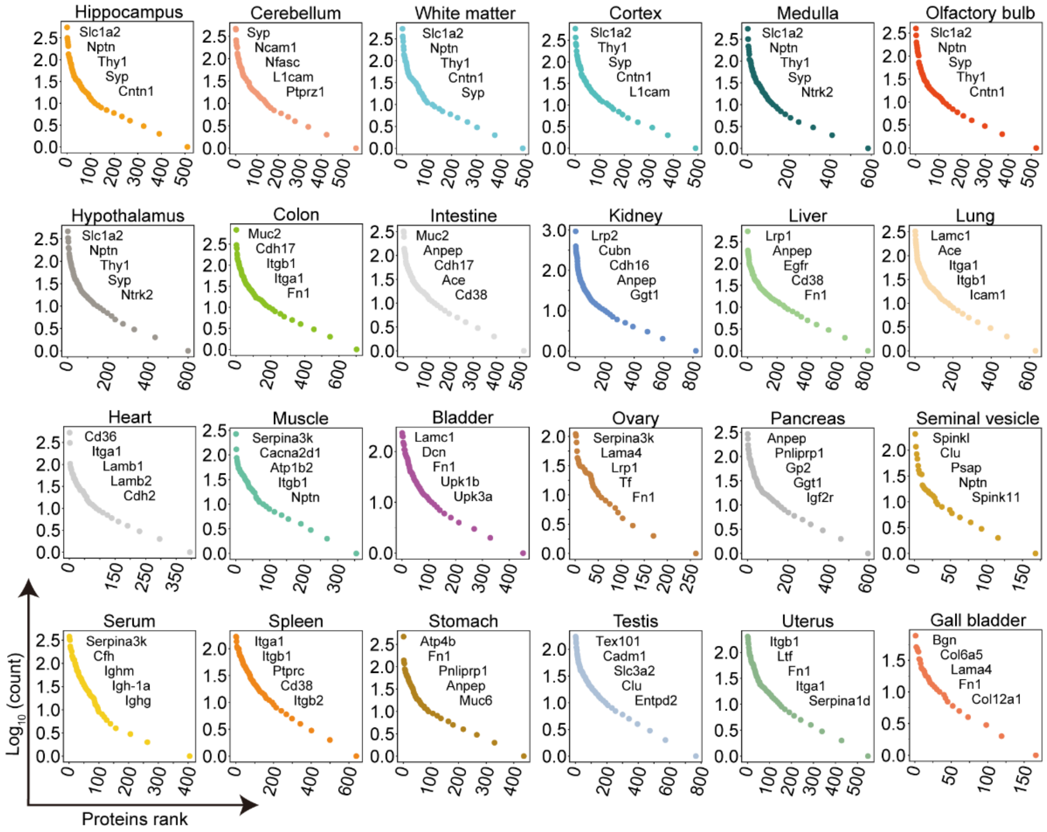


**Figure S7. Comparative clustering of glycopeptides, glycoproteins, and glycans across tissues.** (A) Dynamic range of the intensity-ranked glycoproteins of 24 tissues. Five of the high abundant glycoproteins that relate to the functional specialization of the corresponding tissue are listed in descending order.

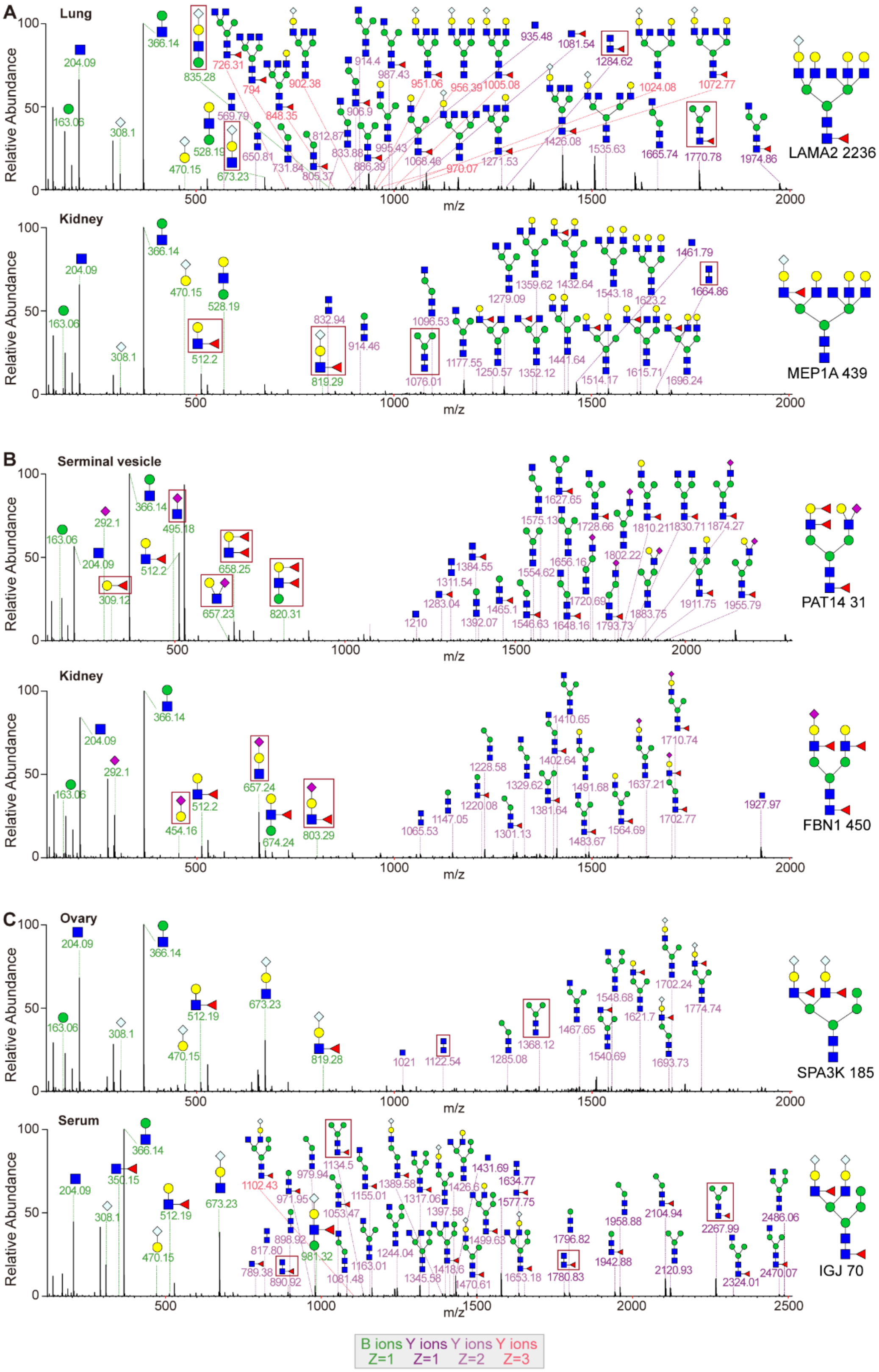


**Figure S8. Representative MS/MS spectra of different glycan isoforms identified in different tissues.**

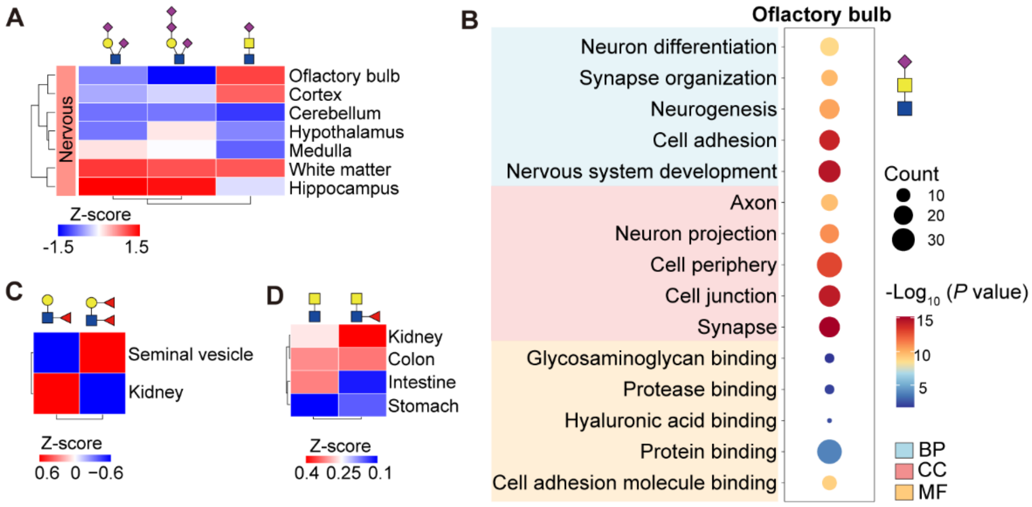


**Figure S9. Structural diversity and tissue-specific distribution of functionally enriched glycan epitopes.** (A) Heatmap showing distribution of sialylated *N*-glycans across seven brain regions. (B) Gene Ontology (GO) enrichment of glycoproteins modified by sialylated LacdiNAc glycans in the olfactory bulb. (C) Distribution of Lewis$^{x/a}$ and Lewis$^{y/b}$ epitopes in the seminal vesicle and kidney. (D) Distribution of LacdiNAc and fucosylated LacdiNAc glycans in the kidney, colon, intestine, and stomach.

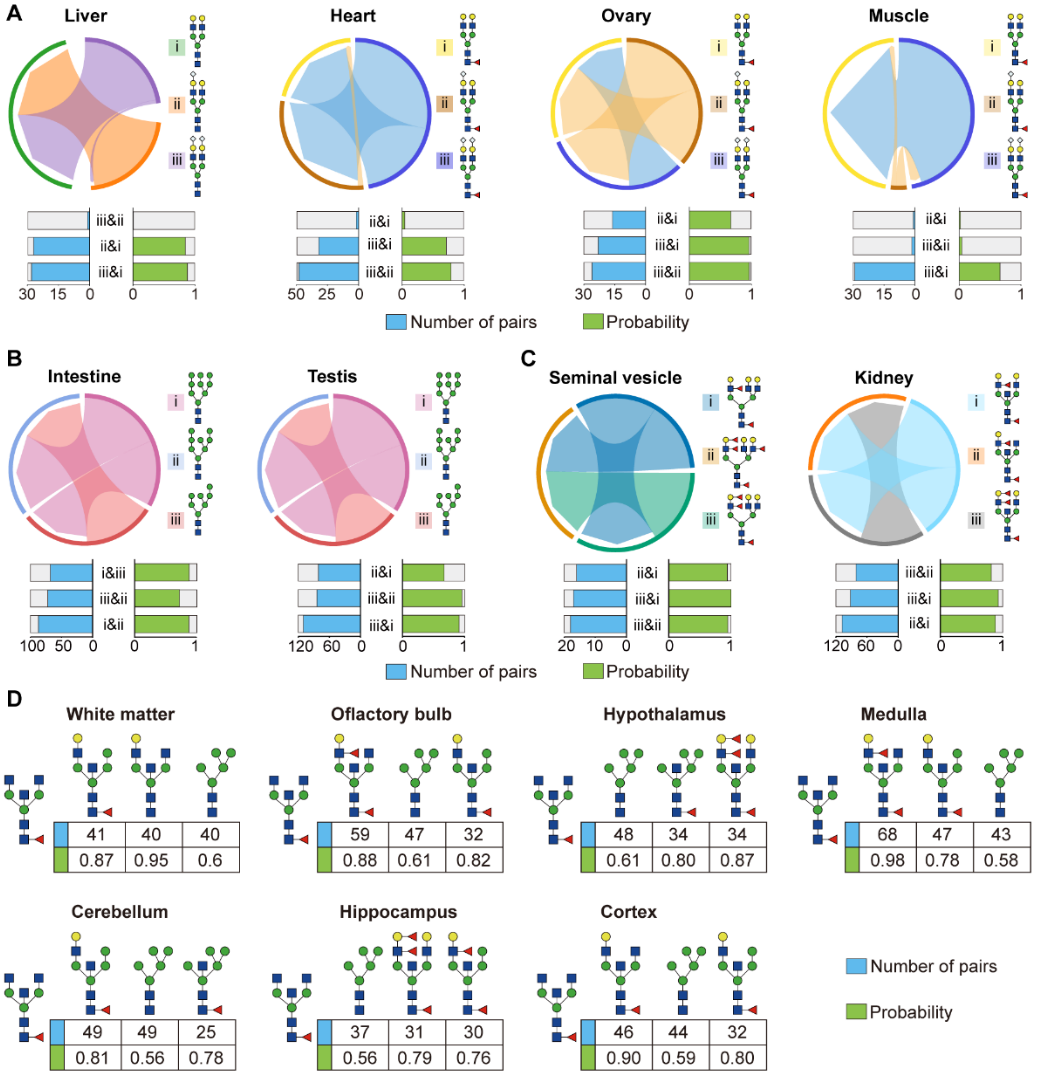


**Figure S10. Representative examples of glycan co-occurrence patterns at single glycosites.** (A)The triplet co-occurrence of sialylated glycans in liver, heart, ovary, and muscle, showing high conditional probabilities. (B) Co-occurrence of oligo-mannose structures (e.g., Man7–Man9) in intestine and testis. (C) Fucosylation-dependent co-occurrence patterns in kidney and seminal vesicle. (D) Co-occurrence patterns of glycans in brain regions.

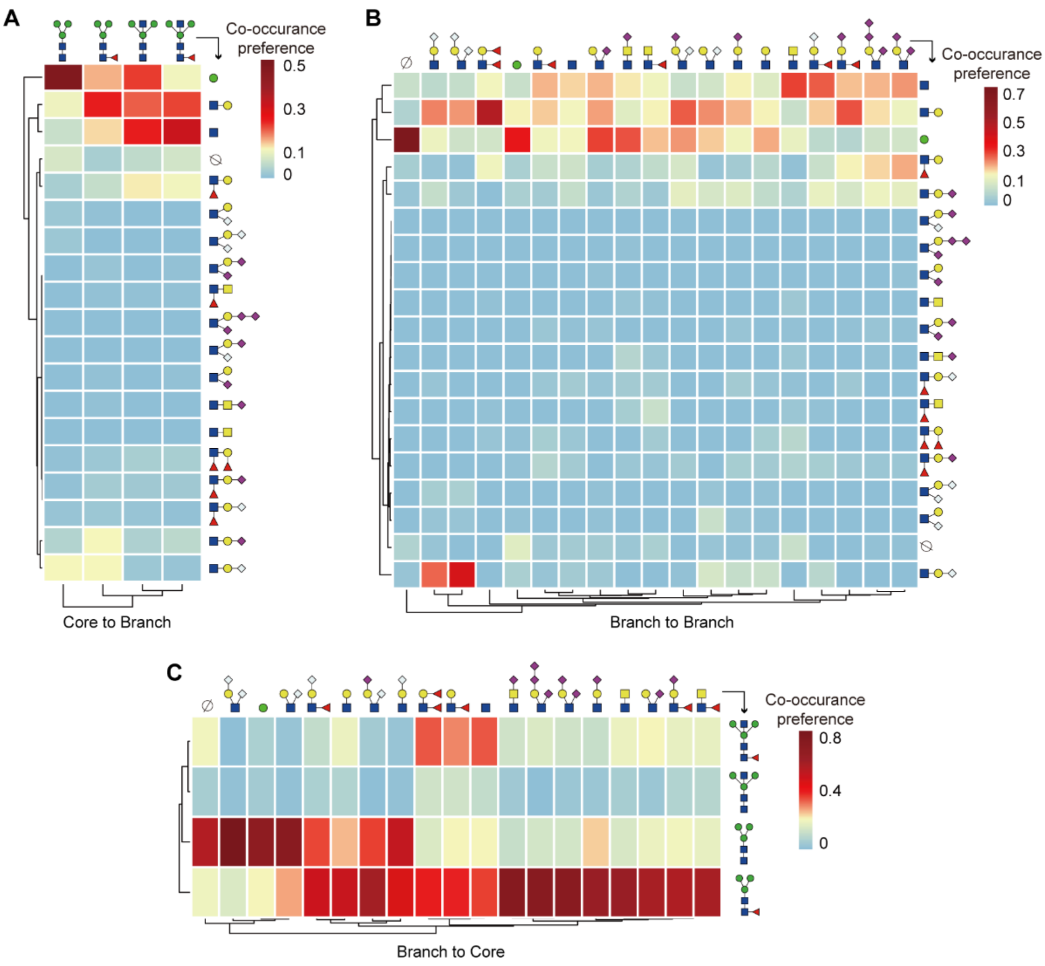


**Figure S11. Heatmap showing the co-occurrence preferences across modular structural features of *N*-glycans.** (A) Core-to-branch associations depict the likelihood of specific branch structures occurring alongside distinct core structures. (B) Branch-to-branch associations show how frequently different branch structures co-occur on the same glycan. (C) Branch-to-core associations indicate the probability of specific core structures appearing with given branch structures. Related to Figure 7C.
